# Genetic determinism of phage-bacteria coevolution in natural populations

**DOI:** 10.1101/2021.05.05.442762

**Authors:** Damien Piel, Maxime Bruto, Yannick Labreuche, Francois Blanquart, Sabine Chenivesse, Sophie Lepanse, Adèle James, Rubén Barcia-Cruz, Javier Dubert, Bruno Petton, Erica Lieberman, K. Mathias Wegner, Fatima A. Hussain, Kathryn M. Kauffman, Martin F. Polz, David Bikard, Sylvain Gandon, Frédérique Le Roux

## Abstract

Coevolution between bacteriophage (or phage) and their bacterial host is thought to be key for the coexistence of these antagonists. Recent studies have revealed the major role of mobile genetic elements in the emergence of phage resistant hosts but how phage escape these defenses in the wild remained to be explored. Here we show a striking parallel in phage evolving counter defenses to host defenses in natural population. We established a large collection of phages and their bacterial hosts and we explored the genetic structure of their interaction. We find that clearly delineated genomic clusters of phage are specific for distinct clades within a bacterial species, *Vibrio crassostreae*, yet while all phages can adsorb, only a subset of hosts are killed due to intracellular defense mechanisms. Host genomes contain multiple mobile defense genes and susceptibility to phage is negatively correlated with genome size. Phages also display extensive gene content variation, but their genome size remains conserved. We show that this gene content variation in hosts and phage is due to rapid turnover of genes involved in defense and escape, and that by exchanging anti-defense genes, phages irreversibly switch host. This could be indicative of co-evolution following the matching-allele-model of specificity and the spatial and temporal variability of phage infectivity further suggests that negative-frequency dependent selection drives phage-vibrio coevolutionary dynamics. We propose a “pan-escape system” that can be shared among phages by homologous recombination within a population that infects a bacterial host.

## MAIN TEXT

The ongoing battle between marine bacteriophages (phages) and their bacterial hosts is probably billions of years old and involves an arsenal of defense and counter-defense systems^1,2^. This arms race is fueled by the underlying coevolutionary dynamics going on at different steps of the infection process^3–5^. Viral infections requires the phage to adsorb to specific receptors at the bacteria cell surface and then bypass intracellular host defenses^6–8^. Recent work on marine *Vibrionaceae* (herein named vibrios) highlighted that phage defense genes turn over exceedingly fast, differentiating clonal isolates, while phage receptors can be highly monomorphic across diverse members of a species, likely due to recent positive selection effecting gene specific sweeps^9^. As a result, phage can enter much more diverse hosts than they can kill. Intracellular anti-phage defense systems are largely encoded by complex, chromosomally inserted mobile genetic elements (MGEs), and single bacterial strain can encode numerous anti-phage systems, suggesting that a wide variety of phages select for multiple resistance systems and/or more than one system is necessary to efficiently prevent infection^10,11^. Bernheim and Sorek recently proposed the ‘pan-immune system’ model which suggests that, although a single bacterial strain cannot carry all possible defense systems because of fitness costs, horizontal gene transfer can allow access to immune defense mechanisms encoded by closely related strains^12^. As phages are thought to rapidly evolve counter-defenses to thrive in the environment, it is expected that the diversity of escape mechanisms within closely related phages mirror the host pan-immune system^12^. However, the counter-defense by phages has been insufficiently studied. While bacteria can considerably expand their genome size, in phage, the size of the capsid also constrains the genome size, likely limiting the number of escape mechanisms a phage can encode. An open question is, therefore, how phage populations counter the numerous defense systems in their hosts and how this influences phage specificity in the wild.

Here we explore the dynamics of phage-bacteria coevolution by focusing on the genetic determinisms of phage-host interactions in natural populations. We combined cultivation, genome sequencing and molecular genetics to analyze a large collection of sympatric and allopatric environmental vibrios and their phages as a model system. We show that genome size, MGE and phage defense element correlate with host resistance to phages. In contrast, phage genome size is conserved among closely related phages, but variable gene content is nonetheless extensive suggesting a role in escaping the host defenses. We demonstrate this in several phages and show that by exchanging anti-defense genes, phages irreversibly switch host.

### Oysters and their vibrio pathogens as a model system

Oysters affected by the Pacific oyster mortality syndrome are infected by diverse virulent strains of *Vibrio crassostreae* that rise to similar abundances in diseased animals^13,14,15,16^. To study the dynamics of *V. crassostreae* at fine temporal resolution, we sampled vibrios from a pool of five juvenile oysters deployed in an oyster farm (Bay of Brest, France) and from the surrounding seawater on 57 dates over five months (each Monday, Wednesday and Friday from May 3 to September 11, 2017) (see Methods). Roughly 48 colonies were picked from *Vibrio* selective plates and screened by PCR targeting the *r5.2* gene, which was previously identified as a *V. crassostreae*-specific marker^14^. Sequencing of the *gyrB* gene confirmed that 195 isolates were *V. crassostreae* (Table S1). Seawater temperature reached 16°C on the 22^nd^ of May, a previously observed threshold for oyster mortalities^17^, and mortalities began on the 29^th^ of May and persisted until the 25^th^ of August (Fig.S1a). *V. crassostreae* occurred only during the disease outbreak with frequencies varying from 0–16% for seawater and 0–58% for oysters (Fig. S1a), consistent with the previously determined increased prevalence of this species in diseased oysters^13^. The quantification of *V. crassostreae* DNA by qPCR further revealed an equal distribution between seawater size fractions (Fig. S1b).

In order to establish a large collection of phages infecting *V. crassostreae*, we combined a sympatric and allopatric sampling strategy. First, we used the 195 *V. crassostreae* strains as “bait” to isolate phages from 20mL-seawater equivalents of viral concentrate (1,000X) or oyster tissues (0.1mg) collected on the same day (Fig. S2). Phage infection was assessed by plaque formation in soft agar overlays of host lawns mixed with a viral source. This approach yielded 45 phages from 18 of 195 tested hosts (9.2%). Second, 90 *V. crassostreae* isolated from 12 June–28 July were screened for phages with: (1) ten pooled seawater viral concentrates from five consecutive dates, and (2) twenty time-shift combinations of single seawater viral concentrates. Each approach resulted in the isolation of 21 and 177 additional phages from 5/90 (5%) and 38/90 (42%) plaque-positive hosts, respectively, in total resulting in a collection of 243 phages from Brest (Table S2). Finally, to better understand the spatial variability of phage infectivity, this collection was complemented with 51 bacteria and 31 phages previously isolated in Sylt (Germany, 2016), where the oyster beds have not yet suffered from *V. crassostreae*-related disease outbreaks^15^.

### Phage-bacteria infection network revealed sparse but modular interaction

To investigate phage host-range, phages isolated in Brest (n=243) and Sylt (n=31) were tested against *V. crassostreae* isolates from the time series (n=117), previously sampled in Brest (n=34) or in Sylt (n=51) and representative members of other *Vibrio* species (n=97), summing to 299 potential hosts. Interactions were assessed by drop-spotting viral lysates onto host lawns to test for plaque formation in an all-by-all assay. To prevent “lysis from without” sometimes observed with high phage concentrations^18^, all phages were normalized to 10^3^ PFU/drop using the original host of isolation. Clearing on plates was assessed after 48 hours. Of the 81,926 tested interactions, only 1,861 were positive (2.2%, Fig. S3). Most phages were specific to *V. crassostreae* with only 14 phages infecting member(s) of other *Vibrio* species. Among these, phage 6E35.1 showed the broadest host range with 26 sensitive *V. crassostreae* isolates from Brest or Sylt and eight strains from other species. Focusing on the *V. crassostreae* species, the matrix suggested that killing tends to occur between subsets of hosts and phages (Fig. S3). However, due to our experimental design, the host-range analysis may be confounded by clonal isolates from the same samples. We considered as potential clones, vibrio isolates with 100% *gyrB* sequence identities and identical patterns of susceptibility in the cross-infection matrix assays. Removal or potential clones resulted in 157 strains, 90 from the time series, and 34 and 33 previously sampled in Brest and Sylt, respectively (Table S3). Phages showing identical patterns of infectivity were also considered clonal, with 76 phages selected as representative of each clone. We found that a minimum set of 24 out of 76 phages was sufficient to kill a maximum of 107 out of 157 (68%) *V. crassostreae* isolates, with a mean infectivity of six hosts per phage.

We sought to explore how the phylogenetic diversity shapes the structure of *V. crassostreae*-phage interaction. We assembled the genome sequences of the 157 isolates, with the number of contigs ranging from 16–824 (Table S3). We observed that the *V. crassostreae* core genome phylogeny formed eight tight clades (V1 to V8) but with different phylogenetic depth (Fig. 1a). Within the less diverse clades, the median single nucleotide polymorphism (SNP) were 900 (clade V1), 1650 (clade V2), 2183 (clade V3), 104 (clade V4) and 2335 (clade V5). The most closely related genomes were separated by 16 single nucleotide polymorphisms (SNP) and considerable gene content variation (Fig. S4), confirming the non-clonality of the isolates. We next characterized the diversity of the *V. crassostreae* infecting phages. Electron microscopy revealed that all phages belong to the *Caudovirales*, with 32, 15, and 29 of 76 being podoviruses, myoviruses, and siphoviruses, respectively (Fig. S5-7). Genome sequencing of these double-stranded DNA viruses revealed that many phages share common genes, and that the genomes overall form distinct genomic clusters using the prefuse force directed layout implemented in Cytoscape^19^ (Fig 1b, Fig. S8).

**Figure 1.**
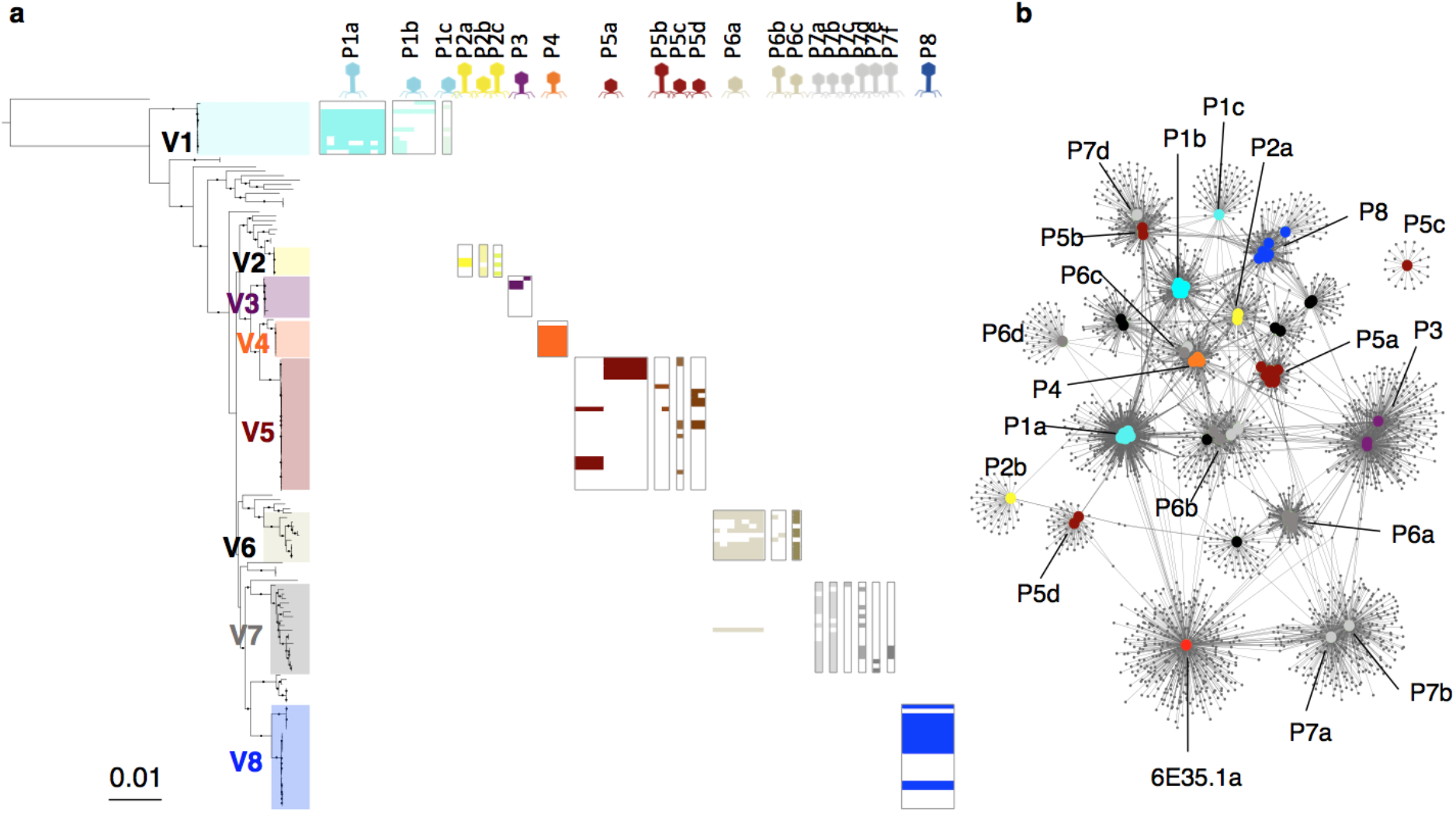
The modularity of the phage-vibrio infection network involves phylogenetic clade within *V. crassostreae* and genomic cluster of phages. **a,** Host range matrix for assay of genome-sequenced phages on genome-sequenced hosts. Rows represent *Vibrio* strains ordered by a Maximum likelihood core genome phylogeny of 157 *V. crassostreae* isolates and *V. gigantis* strain 43_P_281 as an out-group (2498 genes). Clades (V1 to V8) are labeled with different colors. Columns represent phages (n=76) ordered by genomic clusters as defined in b. Vibrio killing by each phage is represented by colored squares. Phage morphotypes are indicated by specific icons for siphoviruses (long tail), myoviruses (medium tail) and podoviruses (short tail). **b,** Phage genome forms clusters. The network was integrated with 2,486 genes family from 76 phages and revealed clustering of phages with genomes (large circles) linked by common genes (small grey circles), %id aa>30% and >80% coverage. The color of each phage genome refers to the clade assignment of the host they kill.

We thus considered the cross-test matrix in light of vibrio core genome phylogeny and phage clustering (Fig. 1a). This revealed that each cluster of phages specifically kills a single clade of *V. crassostreae*. The only exception was the phages from cluster P6a that infect, in addition to strains from clade V6, a single strain from clade V7. This strain was otherwise resistant to phages belonging to cluster P7. Vibrios from a specific clade could be infected by more than one cluster, e.g., vibrios from V1 were killed by phages from cluster P1a, P1b and P1c. We further asked whether the observed specificity of a phage cluster for a vibrio clade results from adsorption variation or intracellular defenses. We show that a representative phage of each of the 16 clusters was only able to adsorb to bacteria from its specific clade of vibrios, with the exception of a phage from cluster P5b that adsorbs to vibrios from clade V5 and V6 (Fig. 2; Fig S9). However, within the same vibrio clade, phages adsorb similarly to all tested vibrios regardless of the production of progeny and cell lysis. Hence clade-specific receptor(s) and cluster-specific receptor binding protein(s) appear to constitute a first level of specificity while within each clade, intracellular mechanisms likely result in a narrower range of vibrio strains that are killed.

**Figure 2.**
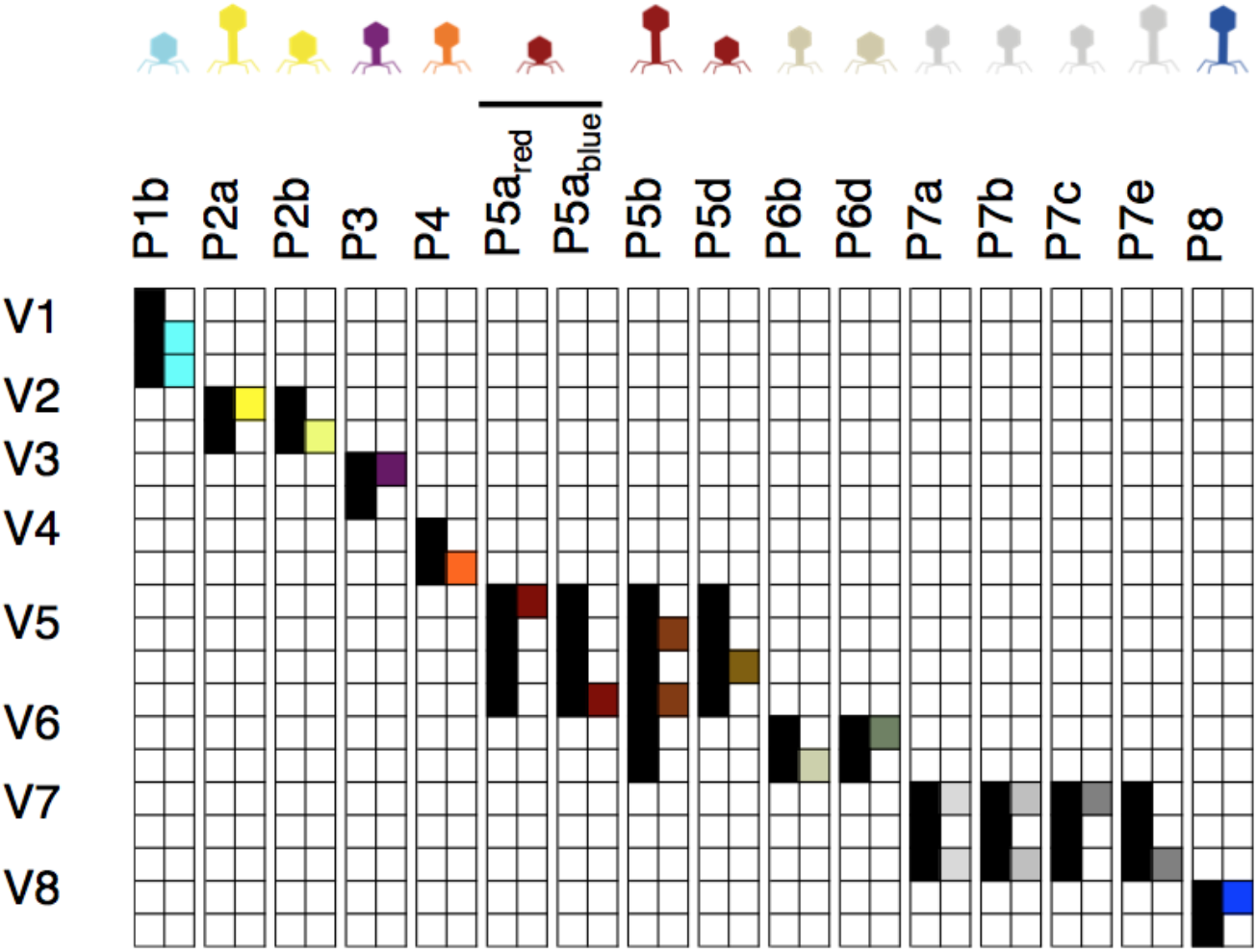
Graphic summary of adsorption and killing assays. Bacteria and phage belonging to a specific cluster are arrayed in rows and columns, respectively. Positive and negative adsorptions are represented by black and white squares respectively. Positive and negative killings are represented by colored and white squares respectively. For phage cluster P5a, two phages (P5a_red_ and P5a_blue_) that differ in their host range are shown. All 320 adsorption assays were performed twice (Fig. S9).

### Flexible genomes of both hosts and phages can be extensive

We observed that *V. crassostreae* genome sequences showed large size variation (Table S3). Strains from clade V1, V6 and V7 had the smallest genomes (medium size 5-5.2 Mbp) and strains from clades V2, V3, V4, V5 and V8 the largest genomes (5.4-5.8 Mbp) (Fig. 3a). The larger genomes contain a higher number of genes encoding for integrases and partitions systems, indicative of a higher number of MGEs. This might allow vibrios to acquire defense elements, as illustrated by a higher frequency of restriction-modification (RM) systems and other known phage defense systems in larger genomes. Accordingly, we observed a negative correlation between genome size and the number of phages able to kill the host (Spearman’s rank correlation of phylogenetic independent contrasts, rho = −0.232, p-value = 0.004). Hence, our results are consistent with a major role of MGEs in the emergence of phage resistance among closely-related strains as well as with the observation that the majority of the pan genome among closely related strains consists of MGEs harboring phage defense genes ^9,12^.

**Figure 3.**
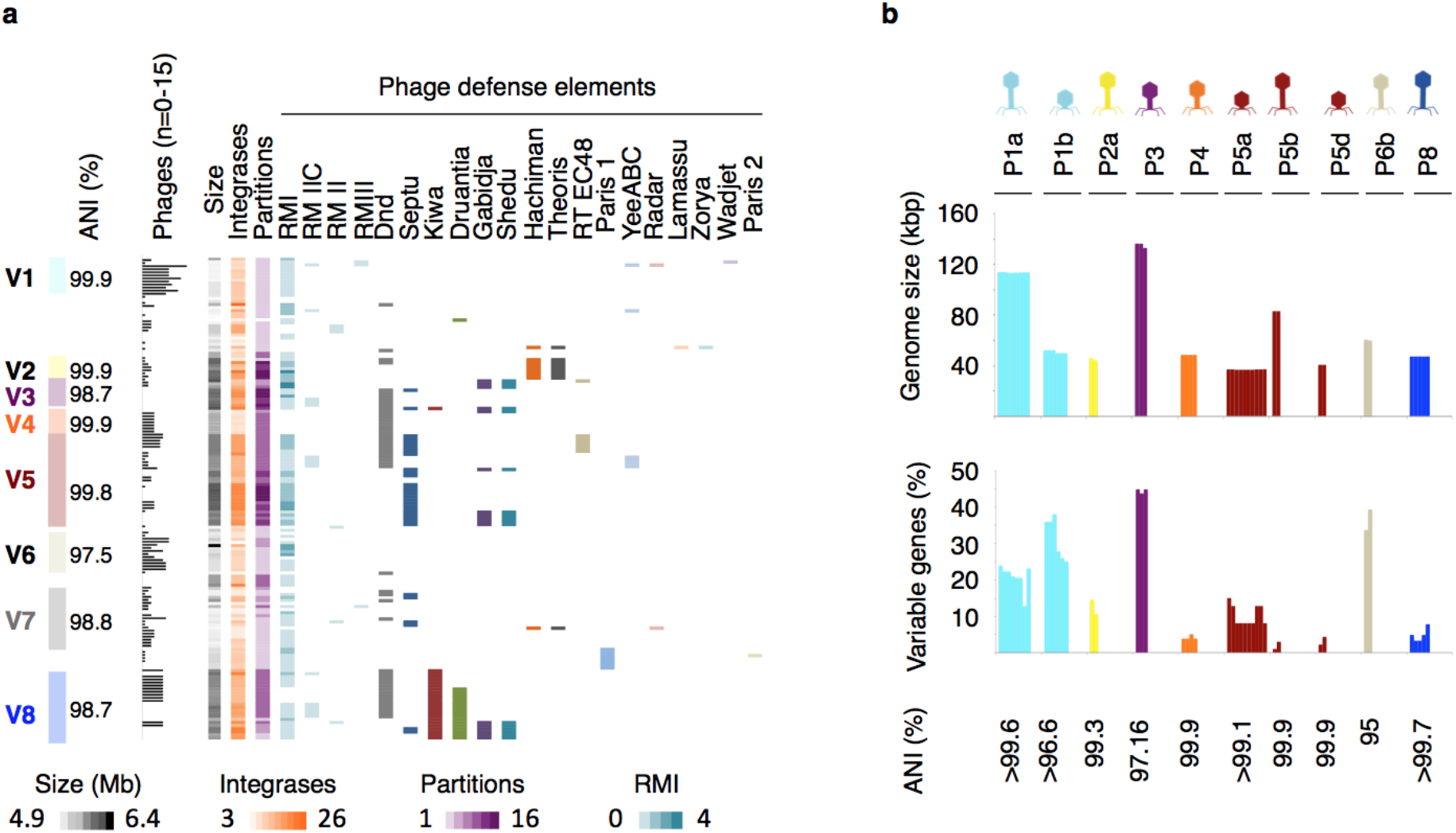
Bacteria and phage flexible genome diversity is indicative of host defense and phage anti-defense interplay. **a,** *V. crassostreae* core genome phylogeny forms clades (V1 to V8 Fig. 1) with specific properties. The ANI value between strains within each clade is indicated. The bar graph indicates the number of phages out of 76 that kill each strain. Columns indicate the genome size (grey); the number of integrases (orange); the number of partition systems (purple); the number of restriction modification systems (blue); the presence of phage defense elements describes by Wang et al., Gao et al., Millman et all, Doron et al., Rousset et al.^10,11,22,24,48^ **b**, intra-cluster comparative genomic indicating the genome size, the percentage of variable genes and the ANI value based on the core genes. Phage morphotypes are indicated on the top by specific icons for siphoviruses (long tail), myoviruses (medium tail) and podoviruses (short tail).

A considerable diversity in genome length (from 21–253 kbp) and gene content (23–407 predicted coding sequences) was also observed for phages (Table S4). Notably the genomes of the podoviruses were found to be significantly smaller than those of the myoviruses and siphoviruses (21–58 kbp, Tukey’s HSD, all p values <0.017) (Fig. S8b) while the broader host range myovirus 6E35.1 has a comparatively larger capsid (Fig. S6) and genome size (253 kb) (Table S4). Intra-cluster genomic comparisons revealed a high conservation of the genome size, a high ANI value of the core genes (>99% in most clusters) but an extensive variation in gene content among phages in some clusters (Fig 3b). Notably a total of 65 putative recombinases (UvsX, Erf, Sak, Sak4, RedB and Gp2.5) were identified in 50 out or 76 phage genomes (Table S4). Altogether our results indicate a recent phage diversification by recombination possibly involving escape mechanisms to host defense, a hypothesis we test further below.

### Within host clades, diverse intracellular mechanisms control phage production

Our genome analysis revealed that vibrios with smaller genomes carry fewer genes with phage defense annotation and tend to be infected by more phages (Fig.3a). We therefore hypothesized that the identification of immunity mechanism should be facilitated in smaller genome hosts that are resistant to phages. In clade V1 (medium genome size 5Mbp), only one strain (7F1_18) out of 12, appeared to be resistant to all siphoviruses from cluster P1a and all podoviruses from cluster P1b. Because cross-infection tests were done at a constant, relatively low phage concentration (10^3^ PFU), we sought to refine estimates of host susceptibility of vibrios from clade V1. First, pairwise interactions were assessed by drop-spotting serial dilution of the phage lysates on host lawns. Second the production of phages was assessed by efficiency of plating (EOP). Combined methods allowed us to classify the strains as “sensitive” or “partially sensitive” if a clearing and viable phage production was obtained at a low or high titer, respectively. The strains were classified as “resistant but impaired” if we observed a turbid clearing zone but no production of viable phages when using high titers. This phenotype may either arise from “lysis from without” (lysis is effected by viral adsorption or extracellular compounds) or abortive infection^18^. Our exploration of host susceptibility at a finer resolution led us to classify the strain 7F1_18 as resistant but impaired to all nine siphoviruses from cluster P1a and a subset of three podoviruses from cluster P1b (named P1b_blue_) (Fig. 4a, b). However, the strain 7F1_18 was partially sensitive to a second subset of P1b phages (named P1b_red_), highlighting diversity among podoviruses from P1b.

**Figure 4.**
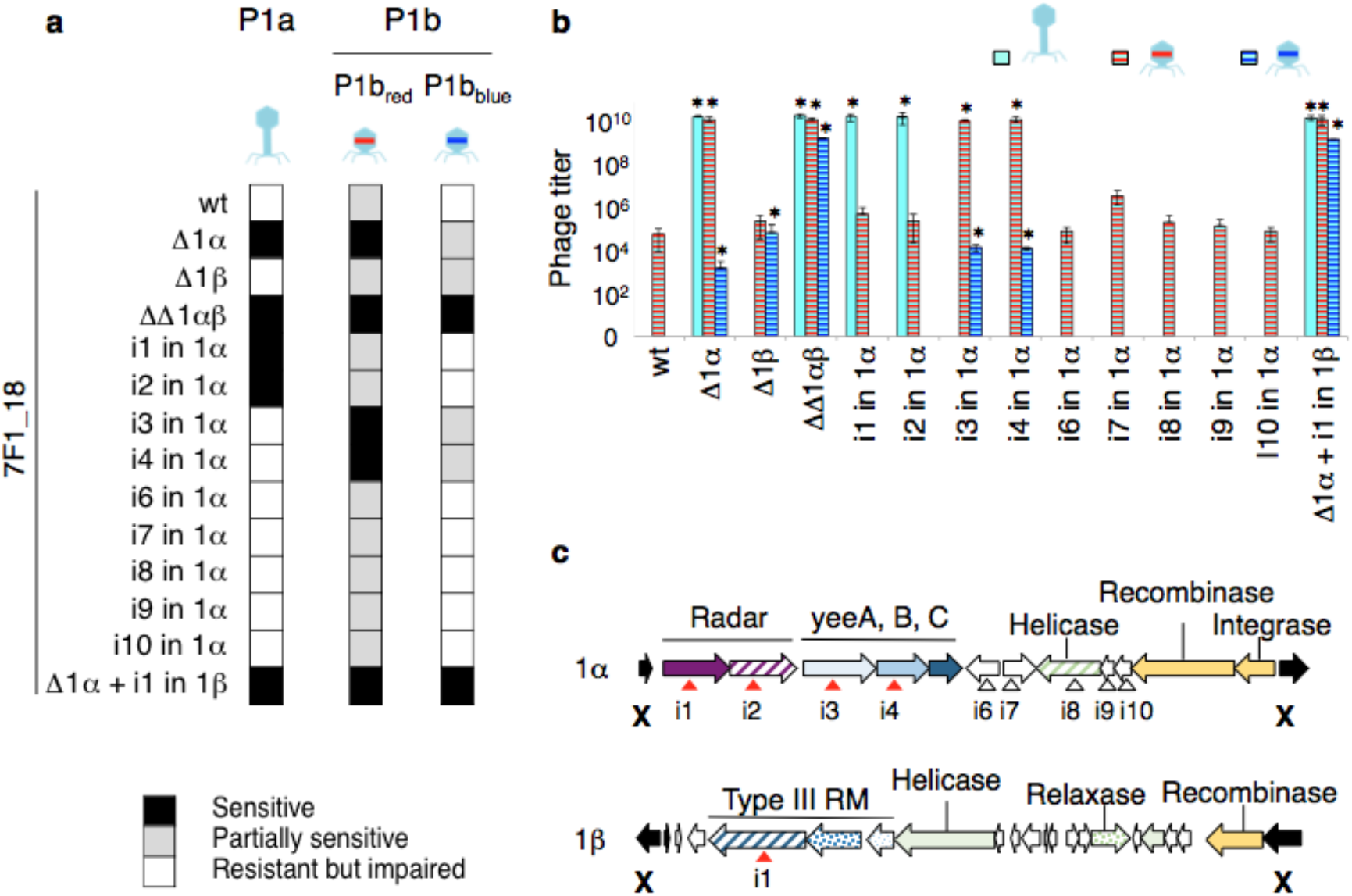
Antiphage elements identified in a strain from *V. crassostreae* clade V1, 7F1_18. **a,** Summary of wild type susceptibility to different phages and changes in susceptibility to the same phages after defense regions deletions or single gene inactivation (see complete results in extended data). **b,** Efficiency of plating (EOP) using representative phages from cluster P1a, P1b_red_ and P1b_blue_ on the wild type (wt) 7F1_18 host strain and its derivatives. All experiments (a and b) were performed twice and showed that phage reproduction strongly depended on the specific combination of phage and gene knock-out (F_26,42_ = 205.20, p < 0.001, * above bars show significant differences for each phage compared to the wt strain). **c,** Gene diagrams of defense regions specific to the 7F1_18. X indicates the 500bp flanking sequence cloned in a suicide plasmid to delete the region by double recombination (see method). Triangles indicate the integration site of a suicide plasmid by single recombination, when conferring a modification in sensitivity in red triangle, when no phenotype in white.

We next explored the genetic determinants of resistance in strain 7F1_18. Genome comparison identified only two genomic regions (1α and 1β) specific to this strain (Fig. 4c). These regions encode for known anti-phage systems. In region 1α, the RADAR defense system consists of an adenosine triphosphatase and a divergent adenosine deaminase that might cause editing-dependent abortive infection in the presence of the phage^11^. The genes *yeeA* and *yeeB*, encode a DNA methylase and a helicase respectively. A type III restriction modification (RM) system^20^ was identified in the region 1β. Genetic knock out of regions 1α and 1β further demonstrated their role in 7F1_18 immunity. The deletion of region 1α was sufficient to restore full sensitivity to all P1a and P1b_red_ phages (Fig. 4a, b and Fig. S10, S11). Single gene inactivation further showed that only Radar was involved in defense to phages P1a while only *yeeAB* mediated resistance to P1b_red_ (Fig. 4a, b and Fig. S10, S11). A subsequent deletion of region 1β or the inactivation of the RM III system was necessary to confer full sensitivity to all P1b_blue_ phages (Fig. 4, Fig. S11). This suggested that P1b_red_ but not P1b_blue_ evolved a RM III escape mechanism. Genome comparison identified two genes encoding for unknown function that are present in all P1b_red_ and absent in all P1b_blue_ phages (Fig. S12). Altogether, our results demonstrate that testing diverse phages will often be necessary to define the role of defense genes and some of them can act additively. Our data also suggested that phages P1b_red_ diversified from P1b_blue_ by acquiring a protection toward a restriction system.

### Bacterial defense and phage anti-defense interplay led to host shift

The examination of the interactions between vibrios from clade V5 and phages from cluster P5a revealed an additional level of modularity (Fig. 5a). Subsets of phages, designated P5a_red_ and P5a_blue_ exclusively killed a subset of vibrios designated V5_red_ and V5_blue_, respectively. We showed above that representatives of P5a_red_ and P5a_blue_ were able to adsorb to all tested V5 strains (Fig. 2). We hypothesized that the specificity of killing depends on the interplay between bacterial defense and phage anti-defense with consequences for phage specificity.

**Figure 5.**
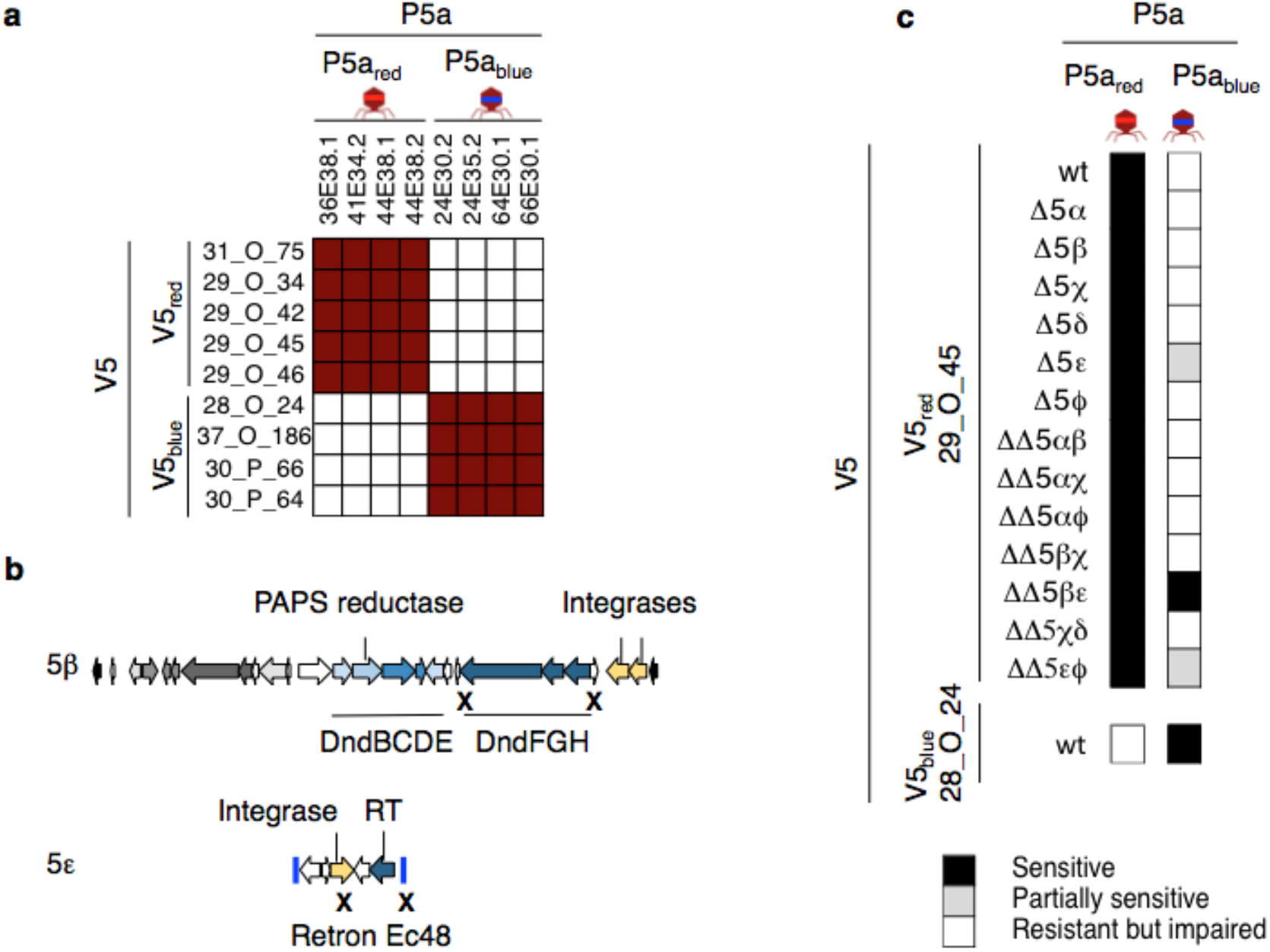
Phage defense elements identified in vibrio V5_red_, and P5_red_. **a,** Modularity of the interactions between vibrios from clade V5 and phage from cluster P5a. **b,** Gene diagrams of two regions specific to the V5_red_ strain 29_0_45 and demonstrated to be involved in resistance to phage P5a_blue_. X indicates the 500bp flanking sequence cloned in a suicide plasmid to delete the region by double recombination (see Methods). Blue lines indicate the end of a contig. **c,** Summary of the changes in susceptibility to phage observed for defense regions deletions (see complete results in extended data). All experiments were performed twice.

Comparative genomics revealed that six genomic regions (5α, 5β, 5χ, 5δ, 5ε, 5ϕ) are found only in all V5_red_ vibrio strains (Fig. S13). This is consistent with the observation that vibrios from clade V5 have larger genomes (medium size 5.7 Mbp) and a higher number of known phage defense elements (Fig. 1). Simultaneous deletions within regions 5β and 5ε resulted in V5_red_ (strain 29_O_45) sensitivity to P5a_blue_ phages (Fig. 5c and Fig. S14). In region 5β (Fig. 5b), the deleted genes (*dndFGH*) are part of the Dnd system, an innate defense system with functional similarity to methylation-based R-M systems^21–23^. DndA-E proteins catalyse phosphorothionate modifications (replacement of oxygen by sulfur in the DNA sugar-phosphate backbone) and the DndFGH proteins use the absence of this modification to identify foreign DNA and cause double-stranded breaks. In region 5ε (Fig. 5b), the two deleted genes encode a reverse transcriptase and a trans-membrane domain protein, homologs of a two-gene phage resistance system, the retron family Ec48, which confers resistance to phage via abortive infection^24^.

EOP allowed higher accuracy in assessment of phage infectivity. P5a_blue_ phages were produced at high levels (10^10^ PFU/ml) in a V5_blue_ host (strain 28_O_24) whereas two orders of magnitude fewer phages were produced in a V5_red_ derivative lacking both Dnd and Ec48 retron defense (ΔDndΔretron) (Fig. 6b). A third deletion in regions 5α, χ, δ or ϕ did not modify this phenotype, suggesting that additional unknown defense mechanism(s) control the full production of phage progeny in V5_red_, strain 29_O_45. No P5a_blue_ phage progeny was produced in the V5_red_ wild type host or a derivative carrying the Ec48 retron and lacking Dnd (ΔDnd). However, P5a_blue_ phages were produced, but at lower titers (10^4^ PFU/ml), in a V5_red_ derivative carrying Dnd and lacking Ec48 (Δretron). In summary, among six genomic regions specific to V5_red_ vibrio, we identified two anti-phage systems that are cumulative, the Ec48 retron being more effective in preventing P5a_blue_ phage production than the Dnd defense system.

**Figure 6.**
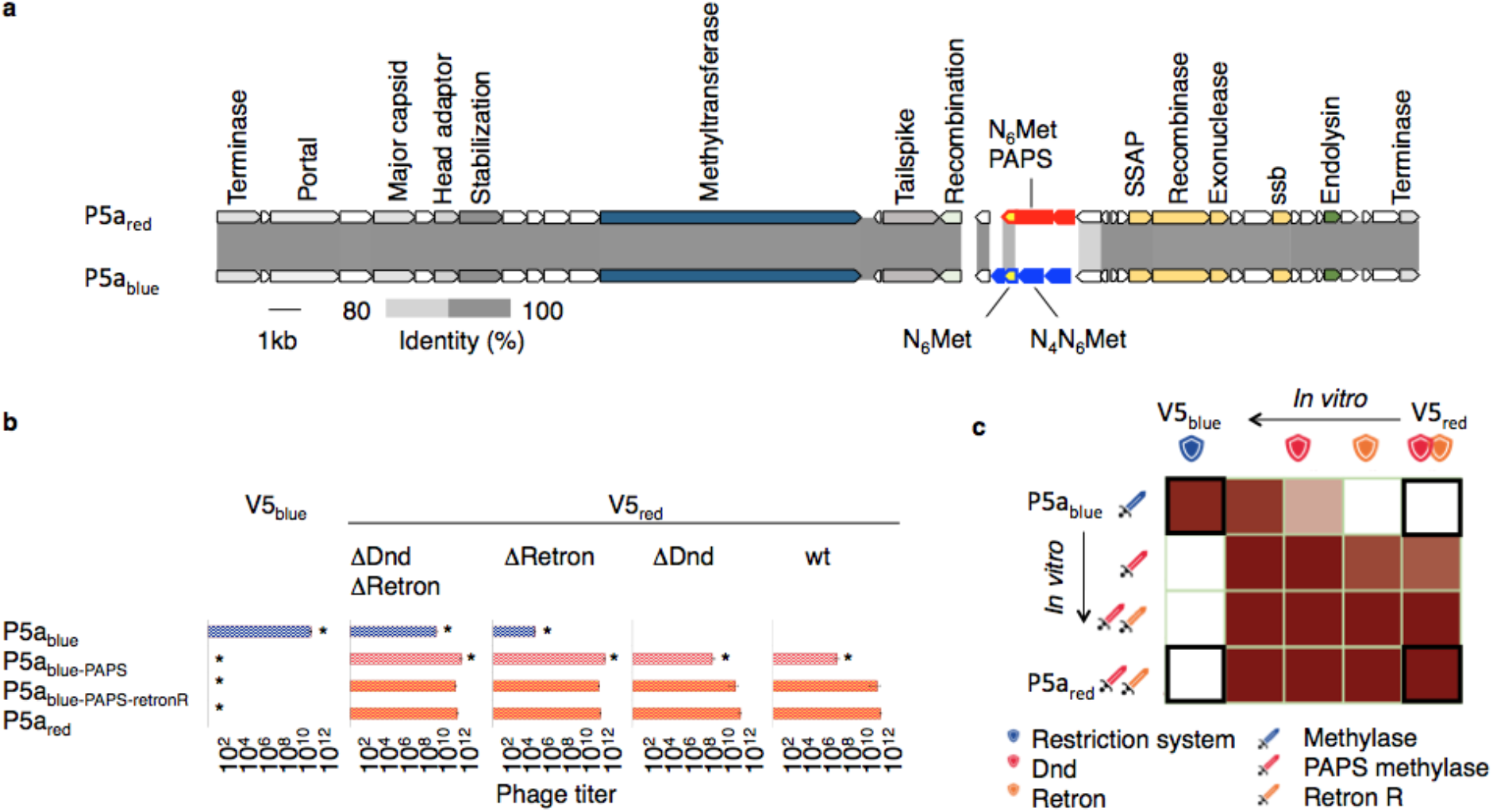
Bacterial defense and phage anti-defense interplay. **a,** Alignment of P5_red_ and P5_blue_ phage genomes showing that gene synteny and content are highly conserved. Only two (red) and four (in blue) genes are found specifically in P5_red_ and P5_blue_ phage. **b,** Number of PFU/ml obtained after vibrio V5_blue_ wild type, V5_red_ wild type and derivatives infection by phage P5_blue_ wild type, P5_red_ wild type and derivatives. All experiments were performed twice and showed that phage reproduction depended on the combination of phage and host derivates (F_12,20_ = 801.49, p < 0.001, asterisk * show significant differences of each phage derivate compared to the V5_red_ wildtype, wt). **c,** Graphic summary of the results. Bold framed indicate the production of phages in combination of phage and host isolated from nature. The other combination results from laboratory manipulation of phage and/or vibrios (in vitro). Colored shields represent anti-phage defense systems acquired by the host (immunity). Colored swords represent the phage anti-defense systems (escape). In the cross matrix, a white square indicates that the phage cannot be produced by the host. Gradients of maroon indicate a production of 10^11^–10^4^ PFU/ml depending on the combination of phage and host tested, as detailed in (a). Exchanging anti-restriction systems allows P5_blue-PAPS_ infection of a new host, V5_red_, but the evolved phage is maladapted to the ancestral host, V5_blue_. When the phage escapes retron defense, of P5_blue-PAPS-retronR_ was similarly infectious to V5_red_ wild type and all derivatives. Indeed P5_blue-PAPS-retronR_ evolved a P5_red_ phenotype.

To understand how P5a_red_ phages evolved to counter vibrio V5_red_ defense systems, Dnd and Ec48 retron, we compared the genomes of podoviruses from cluster P5a. We found only two genes that are specific to P5a_red_ phages (Fig. 6a and S15). Gene p0019 in 44E38.1 encodes a protein of unknown function and gene p0018 encodes a 479 amino acid (aa) protein consisting of two domains: an N-terminal (aa 4–175) phosphoadenosine phosphosulphate reductase (PAPS^25^) domain and a C-terminal (aa 299–470) DNA N-6-adenine-methyltransferase (Dam^26^) domain. A PAPS domain (aa 46–228) is also present in the sulphotransferase encoded by *dndC* in V5_red_ vibrios Dnd defense system. The P5a_red_ Dam domain shares 96% identity with a 178 aa methylase gene (p0019) encoded by P5a_blue_ phage 66E30.1. This suggests a chimeric origin for the p0018 encoded protein, as described for an endonuclease that provides ICP1 phage immunity in *V. cholerae*^27^. Genome comparison also revealed a 5.7kb sequence that diverges between the P5a_blue_ and P5a_red_ phages (Fig. 6a). The region encodes an exonuclease with an RNaseT/DNApolymerase III domain, a single-strand DNA binding protein, two proteins of unknown function and a putative low fidelity single-strand annealing protein (SSAP)-based recombinase system, consisting of two genes similar to λred^28^.

We hypothesized that the incorporation of a PAPS domain by the phage P5a_red_-conferred resistance to the vibrio V5_red_ defense system Dnd. A P5a_blue_ phage was engineered using homologous recombination with a plasmid carrying regions identical to the P5a_red_ and P5a_blue_ phages genome and flanking the two P5a_red_ specific genes (see Methods). This plasmid was transferred by conjugation into a V5_blue_ strain or the V5_red_ derivative Δretron strain. Conjugants were infected by a P5a_blue_ phage (66E30.1) and recombinants were enriched using Δretron as host, because Δretron (i) is partially sensitive to phage P5a_blue_ and therefore allows the production of progeny, (ii) carries the Dnd defense system that might select Dnd-resistant recombinants, and (iii) recombinants might remain sensitive to the retron. We obtained recombinant phages at a high frequency (30%) using Δretron as host for both recombination and selection. All isolated recombinants (designated P5a_blue-PAPS_) were able to infect the V5_red_ derivative Δretron (10^11^ PFU/ml) (Fig. 6b and S16). Thus, the P5a_red_ specific genes encode an anti-Dnd system that is related at least in part to the acquisition of a PAPS domain fused to a methylase.

P5a_blue-PAPS_ phage lost infectivity for V5_blue_ (Fig. 6b, c and S16), demonstrating that the P5a_blue_-specific genes are necessary to infect V5_blue_. Two of the genes encode for methylases (Fig. 6a) in 66E30.1: p0019, annotated as a Dam methylase, and p0020 is a N-4 cytosine-specific and N-6 adenine-specific DNA methylase^29^. These genes likely counteract degradation by V5_blue_ restriction enzyme(s) yet to be identified. Thus by exchanging anti-defense genes, phages irreversibly switch host. This could be indicative of co-evolution following the matching-allele-model of specificity^30^ where an exact genetic match is required for infection.

In an attempt to identify phages that can escape the retron system, we noticed that infection by P5a_blue-PAPS_ resulted in the production of plaques that escape retron immunity (Fig. 6a). Compared to the Δretron vibrio strain, EOPs were, respectively, 10^-5^ and 10^-3^ using V5_red_ and ΔDnd as host. When isolated and further propagated on V5_red_ these escapers showed the same infectivity as P5_red_ (Fig. S16), and are thus likely spontaneous mutants (P5a_blue-PAPS-retronR_). We hypothesized that the 5.7kb sequence that diverges between the P5a_blue_ and P5a_red_ phages isolated in nature is involved in P5a_red_ resistance to retron. Sequencing this region of three laboratory generated retron-escaper phages revealed non-synonymous mutations that distinguished the mutants from the ancestor (Fig. S17), all localized in the exonuclease (p0028 in 66E30.1). Mutations in the exonuclease gene (single mutations, deletions, or integrations) were also observed in 8/10 additional mutants. Spontaneous coliphage mutants have previously been isolated that overcome the defense conferred by Ec48 retron^12^. All mutations abolished the function of RecBCD phage-encoded inhibitors (Gam in λ, gp5.9 in T7), a host complex involved in DNA repair and anti-phage activity. It was proposed that Ec48 “guard” RecBCD and that Ec48 activity is triggered by phage-mediated RecBCD inhibition^12^. None of these inhibitors were identified in the P5a genomes, suggesting that: (i) P5a phages encode non-orthologous protein(s) with similar inhibitory effects on RecBCD; or (ii) retron activity or phage retron escape depends on a mechanism distinct from RecBCD guarding.

### Negative-frequency dependent selection might drive phage-vibrio coevolution in this natural system

Our analyses characterize the genetic basis of antagonistic coevolution between *V. crassostreae* and its phages. While our present sampling density does not allow for an in-depth analysis of coevolutionary dynamics, the spatial and temporal variability of phage infectivity and results from cross-inoculation experiments are consistent with the hypothesis that negative-frequency dependent selection drives phage-vibrio coevolution at the level of phage clusters and vibrio clades. First, across space (Brest *versus* Sylt), phage infectivity is higher on sympatric than on allopatric phage-host combinations (Fig. S18a). This spatial pattern results from the non-overlapping distribution of *Vibrio* clades across locations (Fig. S18b) implying that phages sampled from a given location have a lower chance of finding compatible hosts to infect in allopatry. Second, over time, the mean infectivity peaked for contemporary combinations and declined as phages were inoculated on bacterial strains sampled from more distant sampling dates (“past and future”) ^31–33^ (Fig. S19). The presence of phages of a given clade was significantly associated with the presence of the corresponding *V. crassostreae* clade at that time. This pattern is characteristic of fluctuating selection dynamics, whereby phage populations are maximally adapted to their contemporary bacterial populations^34^. Evidence of modular patterns of specificity at the between-clade level (Fig.1a) and the matching-allele model of specificity within clade V5 (Fig. 6c) further support this hypothesis.

## CONCLUSION

We dissected the genetic mechanisms driving the specificity of the interaction between *V. crassostreae* and their viral predators at different stages of the infection. Phage adsorption matched bacterial clades within the *V. crassostreae* species, suggesting clade-specific receptor(s) and cluster-specific receptor binding protein(s). In the future, the identification of the receptor(s) of each phage cluster should allow us exploring their presence and diversity among the *V. crassostreae* clades and interpretation of how selection acts on these. We can expect that receptor evolution might be constrained due to a role of these surface structures for the fitness of a bacterial clade in the natural environment^35^. Our results are consistent with the previously described major role of MGE in the emergence of phage resistance among closely-related strains and these MGEs constituting the majority of the flexible genome ^9,12^ which is supported here by the variation in vibrio genome size which correlates with resistance, the number of phage defense elements and the identification of diverse anti-phage mechanisms localized in defense genomic regions.

We identify a striking parallel in phage evolving counter defenses to these highly mobile host defenses. First, within a cluster of closely related phages, gene variation can be extensive but the total number of genes per genome is known to be constrained by the capsid side. Accordingly, we identified in podoviruses a mechanism of adaptation by gene exchange rather than gene addition. Exploring phages with larger genomes (such as the myovirus) will decipher whether these phages are more permissive to gene acquisition at multiple loci and/or prone to faster coevolutionary dynamics. Second, homologous recombination system such as the SSAP-like recombinases identified in phage P5a, have been identified in numerous vibriophage genomes and have been suggested to play a role in overcoming bacterial anti-phage defenses by allowing survival of recombinant progeny (Kauffman [*co-submitted*] in revision). This leads us to speculate that gene variation in phage mirror the turnover of MGEs encoding for resistance in the hosts and by analogy to the Bernheim and Sorek model ^12^, we propose a “pan-escape system” that can be shared among phages by homologous recombination within a population that infect a bacterial host.

## MATERIAL AND METHODS

### Sampling

Samples were collected from an oyster farm located at the Bay of Brest (Pointe du Château, 48° 20′ 06.19″ N, 4° 19′ 06.37″ W), every Monday, Wednesday and Friday from the 3^rd^ of May to the 11^th^ of September 2017. Specific Pathogen Free (SPF) juvenile oysters^17,36^ were deployed in the field in batches of 100 animals. When the first mortalities were observed in the first batch, another batch of SPF animals was placed in the field, leading to the consecutive deployment of 7 batches from the 26^th^ of April to the 11^th^ of September. Oyster mortalities were recorded on each sampling day. Oysters were always collected after a minimum of 7 days of incubation in the field.

On each sampling date, five living oysters were collected from a batch showing <50% mortalities. The animals were cleaned, shucked, weighed and 2216 Marine Broth (MB) was added (10mg/ml) for homogenization using an ultra-turrax. A volume of 100 μL homogenate was used for vibrio isolation, the remaining volume was centrifuged (10 min, 10,000 rpm), the supernatant filtered through a 0.2 μm filter and stored at 4°C until the phage isolation stage. Two liters of seawater were collected and size fractionated as previously described^13^. Bacterial cells from 0.2 μm filters were suspended in 2 mL MB and 100 μL of this suspension was used for vibrio isolation. The iron chloride flocculation method^37^ was used to generate 1000-fold concentrated viral samples from 2 liters passaged through a 0.2um filter, following the previously described protocol ^38^. Virus-flocculates were suspended in 2mL 0.1M EDTA, 0.2M MgCl2, 0.2M oxalate buffer at pH6 and stored at 4 °C until the phage isolation stage.

### *Vibrio crassostreae* isolation, identification and genome analysis

#### Isolation and identification

Vibrios from seawater or oyster tissues were selected on Thiosulfate-citrate-bile salts-sucrose agar (TCBS). Roughly 48 colonies were randomly picked from each plate and re-isolated once on TCBS, then on 2216 Marine agar (MA). *V. crassostreae* isolates were first identified by PCR using a primer set targeting the *r5.2* gene (previously identified as population specific marker^14^ (Table S5) and colonies as template. PCR positive isolates were grown in MB and stored at −80°C in 10% DMSO. Their taxonomic assignment was further refined by *gyrB* gene sequencing^14^. Bacteria were grown overnight in MB and DNA extracted using an extraction kit (Wizard, Promega) according to the manufacturer’s instructions. The partial *gyrB* gene was amplified using degenerate primers (Table S5), Sanger sequenced (Macrogen) were manually corrected with the chromatogram. Sequences were aligned with Muscle and phylogenetic reconstruction was done with RAxML version 8 GTR model of evolution, a gamma model and default parameters ^39^.

#### Quantification of V. crassostreae from seawater fractions

Quantification of *V. crassostreae* from seawater size fractions (>60μM, 60-1 μM, 5-1 μM and <1 μM) was performed using quantitative PCR (qPCR). DNA was extracted from filters or 1mg of oyster tissues using the Wizard Genomic DNA extraction kit (Promega). All amplification reactions were analysed using a Roche LightCycler 480 Real-Time thermocycler (Genomic platform SBR). The total qPCR reaction volume was 25 μl and consisted of 4 μl DNA (2,5 ng μl^-1^) and 12,5 μl LightCycler 480 SYBR Green I Master mix (Roche) containing 0.2 μM PCR primer (Table S5) (Eurofins SA) with the following program: enzyme activation at 95°C for 10 min, followed by 40 cycles of denaturation (95°C, 10 s), hybridization (60°C, 20 s) and elongation (72°C, 25 s). A subsequent melting temperature curve of the amplicon was performed to verify the specificity of the amplification. Absolute quantification of bacterial DNA copies were estimated by comparing the observed Cq values to a standard curve of the amplification product cloned into the pCR2.1-TOPO vector.

#### Genome sequencing, assembly and annotation

*V. crassostreae* DNAs were sequenced by the Joint Genome Institute using 300 bp library and HiSeq2000 illumina sequencing technology or at the BioMicro Center at the MIT using Nextera FLEX for library and NextSeq 80PE for sequencing. Contigs were assembled de novo using Spades 3.11^40^. Computational prediction of coding sequences and functional assignments were performed using the automated annotation pipeline implemented in the MicroScope platform^41^.

#### Core genome phylogeny

The proteome of each isolates was compared by performing a Blastp all-vs-all. Silix^42^ was used to reconstruct protein families based on 80% of reciprocal length of alignment and an identity of at least 80% for *V. crassostreae*. Protein sequences of each family were first aligned with Muscle, filtered using Gblocks with relaxed parameters^43^ and concatenated. Phylogenetic reconstruction was done using RAxML version 8^39^ on this concatemer using an LG model of evolution, a gamma model and default parameters.

#### Comparative genomics

The ANI-value of genomes was determined using pyani (https://github.com/widdowquinn/pyani). The phylogenetic profile method implemented in the MicroScope platform^41^ was used to identify putative phage resistance genes and regions. To this aim we searched for genes present in all strains resistant to phage (80% identities on 80% coverage) and absent from sensitive strains. The same approach was used to estimate specific genes of *Vibrio* in pairwise genome comparisons.

### Phage isolation, identification and genome analysis

#### Isolation and generation of high titer stocks

We used the methods previously described by Kauffman and coll ^38^. Briefly isolation of phages was performed by directly plating on a bottom agar plate (1.5% agar, in MB) 100 μL of an overnight bacterial culture, 20 μL of seawater flocculate (equivalent to 20 mL of seawater containing viruses) or 20 μL of oyster homogenate and 2.5 ml molten top agar (55 °C, 0.4% agar, in MB) to form host lawns in overlay and allow for plaque formation. After incubation for 48h at room temperature (RT), a maximum of six plaques per morphotype was archived. Plaque plugs were first eluted in 500 μl of MB for 24 h, 0.2-μm filtered to remove bacteria, and re-isolated three times on the sensitive host for purification before storage at 4°C and, after supplementation of 25% glycerol at −80°C. High titer stocks (>10^9^ PFU/ml) were generated by confluent lysis in agar overlays.

#### Electron microscopy

Following concentration on centrifugal filtration devices (Millipore, amicon Ultra centrifugal filter, Ultracel 30K, UFC903024), 20 μl of the phage concentrate were adsorbed for 10 min to a formvar film on a carbon-coated 300 mesh copper grid (FF-300 Cu formvar square mesh Cu, delta microscopy). The adsorbed samples were negatively contrasted with 2% Uranyl acetate (EMS, Hatfield, PA, USA). Imaging was performed using a Jeol JEM-1400 Transmission Electron Microscope equipped with an Orious Gatan camera. at the platform MERIMAGE (Station Biologique de Roscoff, France).

#### DNA extraction, sequencing, assembly and annotation

Phage DNA extractions were performed from high titer suspensions using the MasterPureTM Complete DNA and RNA Purification Kit (Epicentre), according to the manufacturer’s instructions. Alternatively, DNA was extracted following a previously described protocol ^38^. Phage suspensions were concentrated on centrifugal filtration devices (30 kDa Millipore Ultra Centrifugal Filter, Ultracel UFC903024) and washed with 1/100 MB to decrease salt concentration. The concentrates were treated for 30 min at 37°C with 10μL of DNAse (Promega) and 2,5μL of RNAse (Macherey-Nagel) at 1000 unit and 3,5mg/mL, respectively. These nucleases were inactivated by adding EDTA (20 mM, pH8). DNA extraction encompassed a first step of protein lysis (0.02 M EDTA pH 8.0, 0.5 mg/ml proteinase K, 0.5% sodium dodecyl sulfate) for 30 min incubation at 55°C, a phenol chloroform extraction and an ethanol precipitation. DNA was visualized by agarose gel electrophoresis (0.7% agarose, 50 Volt, overnight at 4°C) and quantified using QuBit. Phages were sequenced by the Biomics platform at the Pasteur Institute using NextSeq Illumina technology. The assembly, annotation and comparative analysis were performed as described above for *V. crassostreae* genome.

#### Phage clustering

The phage proteome was used to reconstruct a network showing the shared families using the force-directed layout implemented in Cytoscape^19^

### Host range determination

#### Single-phage-by-single-host host range infection assay

Host range assays were carried out using a robot hosted at EligoBioscience (Paris, France) or manually using an electronic multichannel pipette by spotting 5 μL of the phage suspension normalized at 2×10^5^ PFU/ml (10^3^ PFU/spot) on the agar overlay inoculated with the tested host. Plates were incubated overnight at room temperature and plaque formation was observed after 24 hours. Spot assays were performed in duplicate and positive interactions were confirmed in a third experiment.

#### Classification of host sensitivity

To explore the sensitivity of bacteria, 10-fold serial dilutions of phages (1-10^-7^ PFU) were prepared and 5 μL drop spots of each dilution were pipetted onto bacterial host lawns. For some spot tests, turbid plaques were observed for the highest concentrations of phage lysates. To determine whether the bacterial host was sensitive, partially sensitive or insensitive but impaired, we explored the titer of the phage on a given bacteria compared to the maximum titer observe (i.e. with the host used to produce the phage). A total of 5μL of serial phage dilutions was mixed with 100 μL of an overnight host culture and 2,5 ml top agar to form host lawns in overlay and plaques were counted after 24hours. In sensitive and partially sensitive hosts plaques were obtained using 1-10 and 10^5^-10^6^ PFU respectively. In resistant but impaired host no plaque was observed using up to 10^7^ PFU.

#### Phage adsorption

Phage adsorption experiments were performed as previously described ^44^. Phages were mixed with exponentially growing cells (OD0.3; 10^7^ CFU/mL) at a MOI of 0.01 and incubated at RT without agitation. At 0, 15 and 30 minutes, 250 μL of the culture was transferred in a 1.5 mL tube containing 50 μL of chloroform and centrifuged at 14,000 rpm for 5 min. The supernatant was 10-fold serially diluted and drop spotted onto a fresh lawn of a sensitive host to quantify the remaining free phage particles.

### Molecular microbiology

#### Strains and plasmids

All plasmids and strains used or constructed in the present study are described in Table S6 and S7. *V. crassostreae* isolates were grown in Luria-Bertani (LB), or LB-agar (LBA) +0.5 M NaCl at RT. *Escherichia coli* strains were grown in LB or on LBA at 37°C. Chloramphenicol (5 or 25μg/ml for *V. crassostreae* and *E. coli*, respectively), thymidine (0.3 mM) and diaminopimelate (0.3 mM) were added as supplements when necessary. Induction of the P_BAD_ promoter was achieved by the addition of 0.2% L-arabinose to the growth media, and conversely, was repressed by the addition of 1% D-glucose. Conjugation between *E. coli* and *Vibrio* were performed at 30°C as described previously^45^ with the exception that we used TSB-2 (Tryptic Soy Broth supplemented with 1,5% NaCl) instead of LB for mating and selection.

#### Clonings

All clonings in pSW7848T were performed using herculase II fusion DNA polymerase (Agilent) for PCR amplification and the Gibson Assembly Master Mix (New England Biolabs, NEB) according to the manufacturer instructions. Before cloning in pSW23T, a PCR fragment was amplified using GoTaq DNA polymerase (Promega) and subcloned in a TOPO cloning vector (Invitrogen). The plasmid miniprep was digested with *EcoR1* (NEB) and the insert was cloned in pSW23T. The phage specific region was amplified using the herculase, digested with *Apa1* and *Xba1* and cloned in PSU18T-P_BAD_ instead of the *gfp* gene. All clonings were first confirmed by digesting plasmid minipreps with specific restriction enzymes and second by sequencing the insert (Macrogen).

#### Vibrio mutagenesis

Gene inactivation was performed by cloning an internal region of the target gene in the suicide plasmid pSW23T^46^. After conjugative transfer, selection of the plasmid-borne drug marker (Cm^R^) resulted from integration of pSW23T in the target gene by a single crossing-over. Region deletion was performed by cloning 500bp fragments flanking the region in the pSW7848T suicide plasmid ^45^. This pSW23T derivative vector encodes the *ccdB* toxin gene under the control of an arabinose-inducible and glucose-repressible promoter, *P_BAD_* ^45^. Selection of the plasmid-borne drug marker on Cm and glucose resulted from integration of pSW7848T in the genome. The second recombination leading to pSW7848T elimination was selected and arabinose media.

#### Phage mutagenesis

P5a_blue_ phage was engineered using double crossing over with a plasmid carrying regions of homology (438 and 156 bp) to the phage genome P5a_red_. A 3745bp region of the phage P5_red_ (44E38.1) was amplified by PCR and cloned in a replicative plasmid (P15A *oriV*; CmR) under the control of the conditional *P_BAD_* promotor. Selection of the transformants on Cm + Glucose 1% prevented the expression of toxic phage genes. This plasmid was transferred by conjugation to a V5_blue_ strain (28_O_24). Plate lysates were generated by mixing 500 μl of an overnight culture of the transconjugant with the P5_blue_ phage (66E30.1) and plating in 7.5 ml agar overlay. After the development of a confluent lysis of lawns, the lysate was harvested by addition of 10 mL of MB, shredding of the agar overlay and stored ON at 4°C for diffusion of phage particles. The lysates were next centrifuged, the supernatant filtered through 0.2 μm filter and stored at 4°C. Recombinant phages were enriched by infecting the P5_red_ (29_0_45) derivative Δretron in agar overlays. Recombinant phages were screened by PCR using a primer set targeting the P5_red_ specific gene (PODOV008_V2_p0019 in 44E30.1) and single plaque as template. The recombination was further confirmed by sequencing genes that are polymorphic between between P5_red_ and P5_blue_ phages.

To isolate mutant phages that escape Ec48 defense, P5_red_ or P5_red-PAPS_ phages were plated on V5_red_ wild type or ΔDnd derivative using the double-layer plaque assay. Plaques were obtained only using the P5_red-PAPS_ as viral source and 10 single plaques were picked for reisolation. The region (from p0024 to p0032 in 66E30.1, 5.3kb) that shows polymorphism between the P5_red_ and P5_blue_ phages was PCR amplified sequenced. Reads were aligned to the ancestor genome.

### Time shift analysis

To characterize coevolutionary dynamics between *V. crassostreae* and its phages, we examined how phage infectivity varied with the time shift between *V. crassostreae* and phage isolates ^32,47^. This was done for 1 to 15 *V. crassostreae* colonies per sampling day (median 3 colonies), and 1 to 46 47 phage strains (median 6 strains), for a total of 254,974687 cross-inoculations. Infectivity was a binary variable coding whether the phage can infect or not the bacterial isolate. The mean infectivity as a function of time shift category peaked around the present and declined as phages were inoculated on bacterial strains of the past and the future (Fig. S19a). This pattern is characteristic of fluctuating selection dynamics, whereby phage populations are maximally adapted to their contemporary bacterial populations. We fitted to this pattern a smooth unimodal relationship (proportional to the density of a skew-normal distribution) by least-squares (Fig. S19b).

It is difficult to statistically test for the significance of this pattern, as it emerges from non-independent combinations of *V. crassostreae* and phage strains, and with limited sampling of diverse populations at each time point that can generate spurious temporal fluctuations. As a simple approach, we compared the difference in infectivity of contemporary vs. non-contemporary combinations. Phages could infect contemporary bacteria in 156211/1,843392 combinations (frequency: 0.11105), and non-contemporary bacteria in 12151188/243,14915 combinations (frequency: 0.049524). The difference was significant according to a test based on binomial probabilities (p = 0.0085799 for the null hypothesis that the true frequency is the same for the two groups).

## Acknowledgements

We thank Marie Agnes Petit and Melanie Blokesch for valuable suggestions and Marie Touchon and David Goudenège for assistance for vibrio genome annotation. We thank Zoe Chaplain for the illustrations and help during the time series sampling. We thank the staff of the station Ifremer Argenton and Bouin, the ABIMS (Roscoff) and LABGeM (Evry) platforms for technical assistance. We thank Guy Riddihough from Life Science Editors for help with the Manuscript.

## Author contributions

FLR and DB conceived of the project. FLR wrote the paper with contributions of DP, MB, YL, FB, MW, FAH, KK, MP, DB and SG. DP performed phage-vibrio interaction experiments with assistance from SC, RBC and EL. MB performed the *in silico* analyses with assistance from KK. YL and FLR performed the genetics. SL performed the electronic microscopy analyses. DP, YL, SC, AJ, BP and FLR established the times series sampling. MKW, FLR and JD isolated the phage and vibrio collections from Sylt. FAH and MP performed and funded part of the vibrio sequencing. FB and SG designed, FB performed the time-shift analysis. FLR supervised the project and secured funding.

## Data and materials availability

New genomes used in this work have been deposited under the NCBI BioProject with accession numbers presented in Table S3 and S4. All data, code, and materials are available upon request.

