## Supplementary Figures for "Genetic determinism of phage-bacteria coevolution in natural populations"

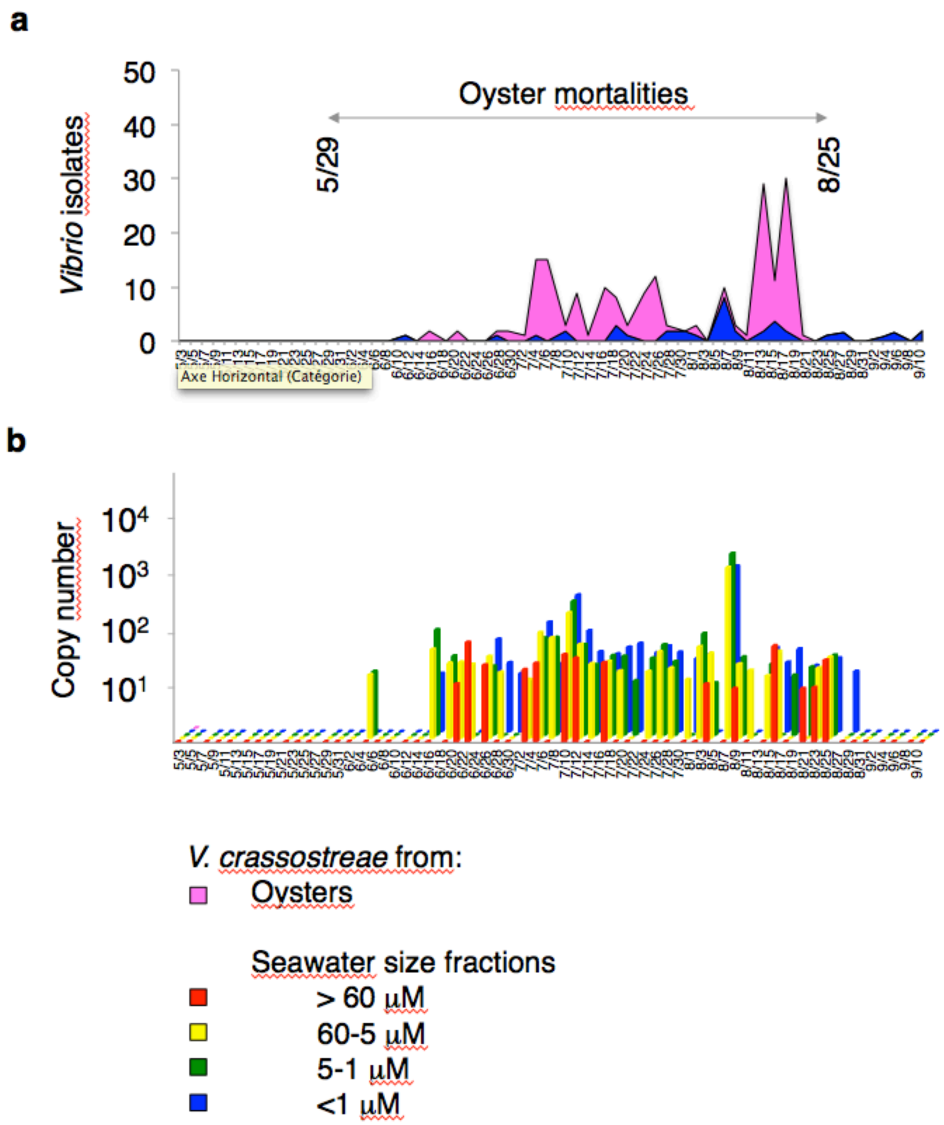


**Figure S1. Dynamics of *V. crassostreae*. a,** On each sampling date (3 May-11 September 2017, x-axis), total vibrios from the seawater (size fraction <1µM, blue) or five oyster tissues (dark pink) were selected on TCBS and genotyped to identify *V. crassostreae* isolates. The y-axis indicates the number of positive isolates out of the randomly picked 48 colonies. The arrow indicates the period of oyster mortalities, i.e. 29 May-25 August. **b,** *Vibrio crassostreae* abundance measured by qPCR was shown to be similar between the different seawater fractions.

| *V. crassostreae* | | Viral sources | Number of assays | Plaque positive hosts | Number of Phages |
|---|---|---|---|---|---|
| **1: Combination from the same day** | | | | | |
| 195 isolates (June 12- September 9) | X | 1 date, seawater<br>1 date, oysters | 195x2=390 | 18 (9.2%) | 45 |
| **2: Pool of viruses from seawater** | | | | | |
| 90 isolates (June 12-July 28) | X | 10 pools of 5 dates seawater | 90x10=900 | 5 (5%) | 21 |
| **3: Time shift combinations** | | | | | |
| 90 isolates (June 12-July 28) | X | 20 dates seawater | 90x20=1800 | 38 (42%) | 177 |

**Figure S2.** The three procedures used to isolate phages against *V. crassostreae* during the time series sampling in Brest.

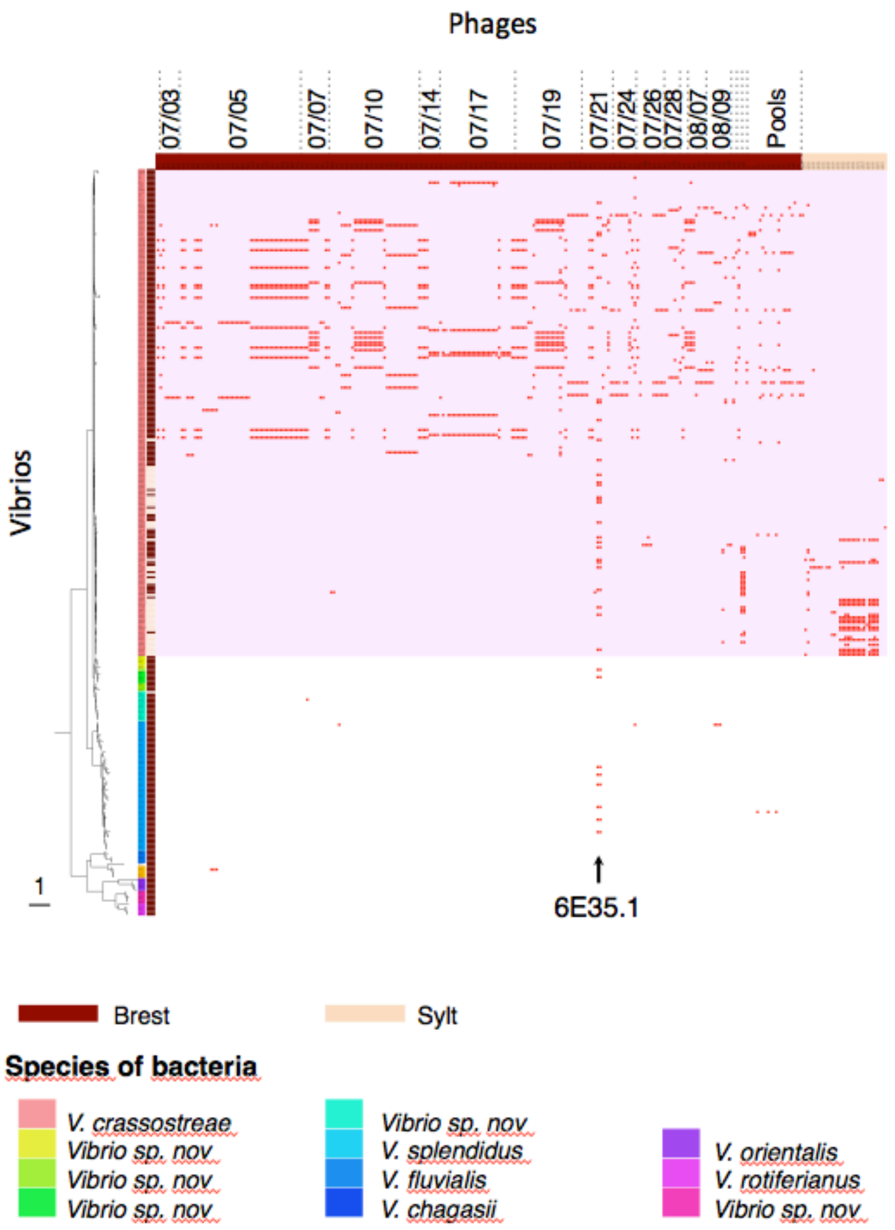


**Figure S3.** Host range matrix for assay of phages on *gyrB*-sequenced hosts. Rows represent *Vibrio* strains (n=299), columns represent phages (n=243 isolated in Brest and ordered by date or n=31 isolated in Sylt), red marks indicate infection of host. Pink shade discriminate *V. crassostreae* isolates from representatives of other species.

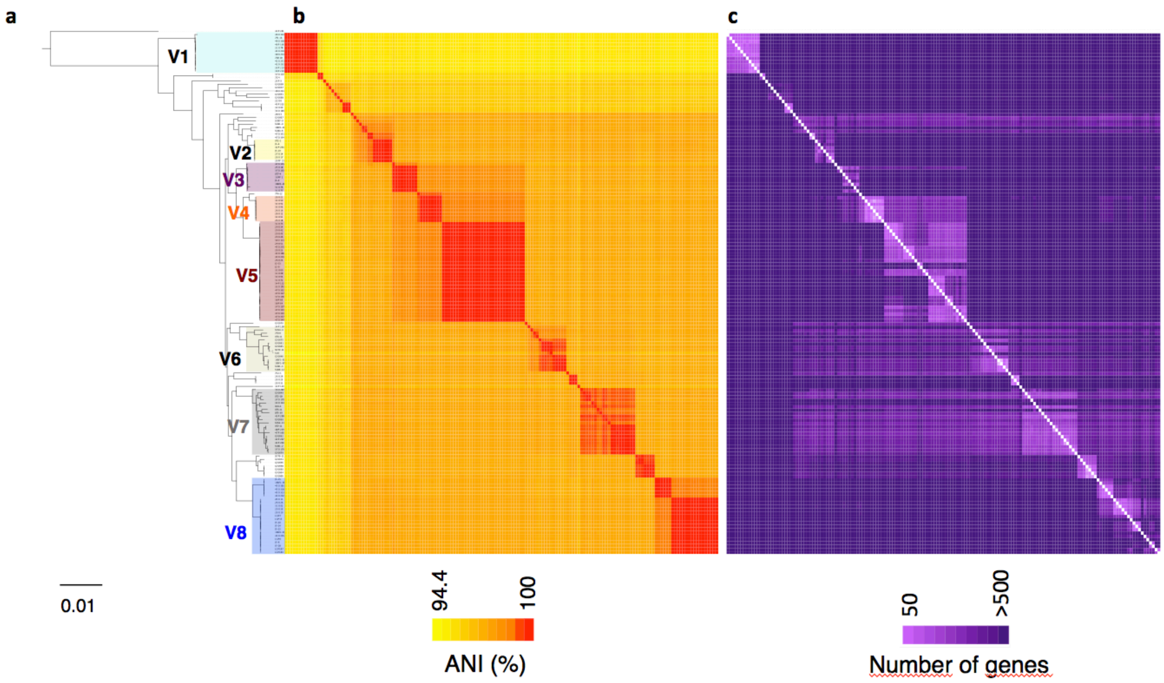


**Figure S4**. **Genome diversity of sequenced *V. crassostreae.* a,** Core genome phylogeny of 157 *V. crassostreae* isolates (2498 genes, Maximum Likehood) and *V. gigantis* strain 43_P_281 as an out-group. **b**, Pairwise ANI values showing vibrio hosts from different clades (colored shade) encompasses nearly clonal strains (clade V1 to V5) or more diverse strains (clade V6, 7 and 8). **c,** accessory genome heatmap.

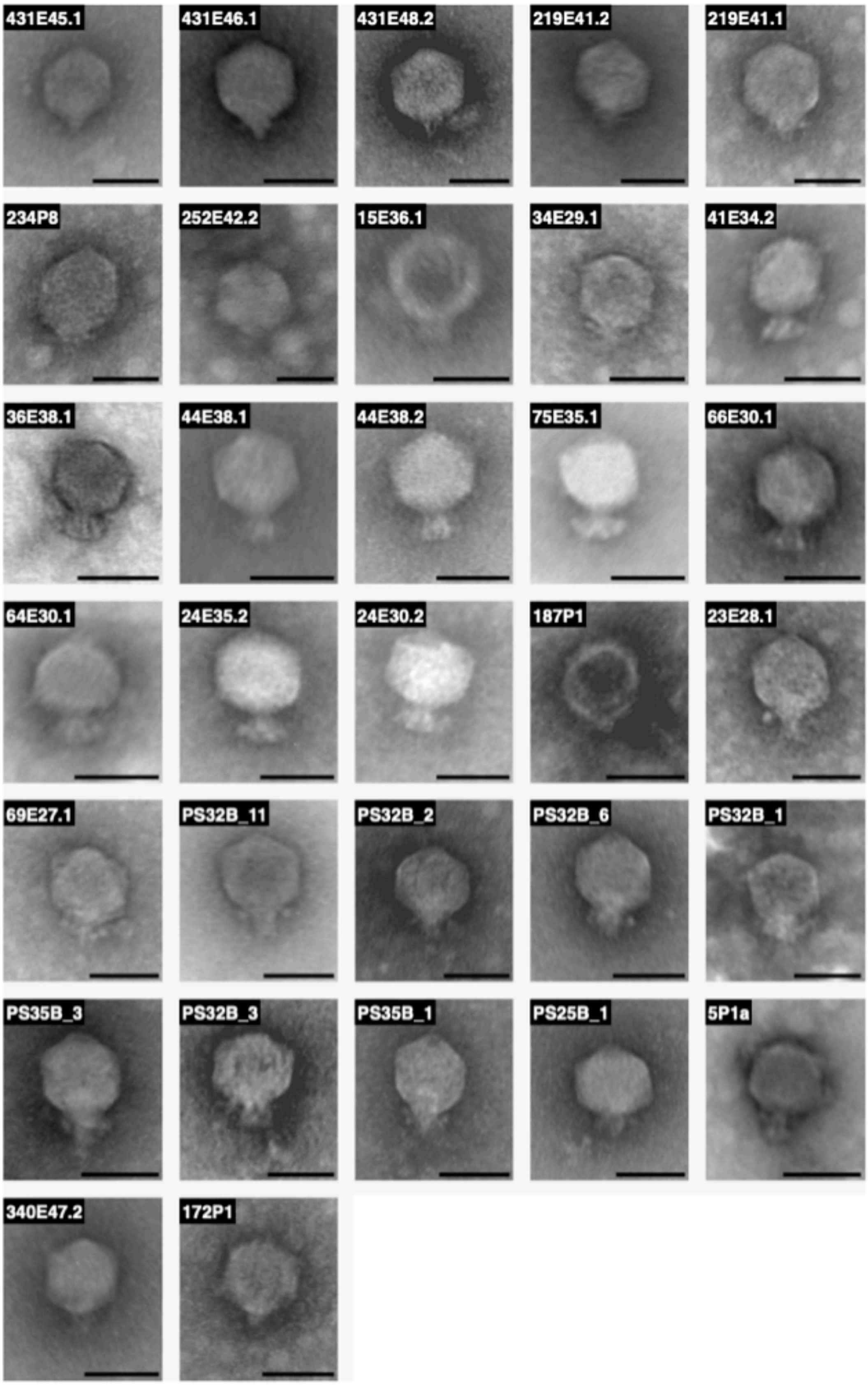


**Figure S5.** Transmission electron micrographs of 32 podovirus. Scale bars represent 50nm

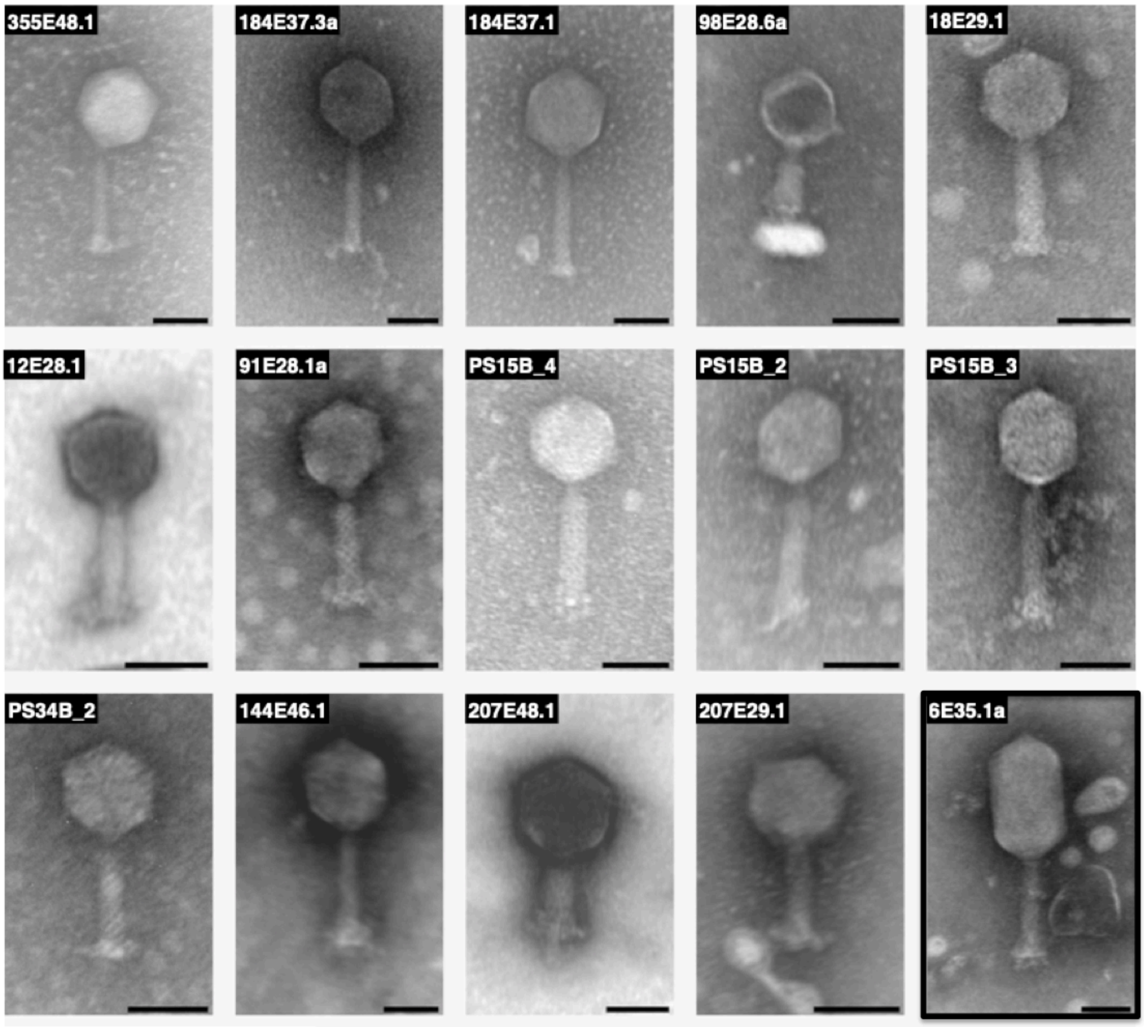


**Figure S6.** Transmission electron micrographs of 15 *Myoviridae.* The broader host range phage 6E35.1a is surrounded in bold. Scale bars represent 50nm

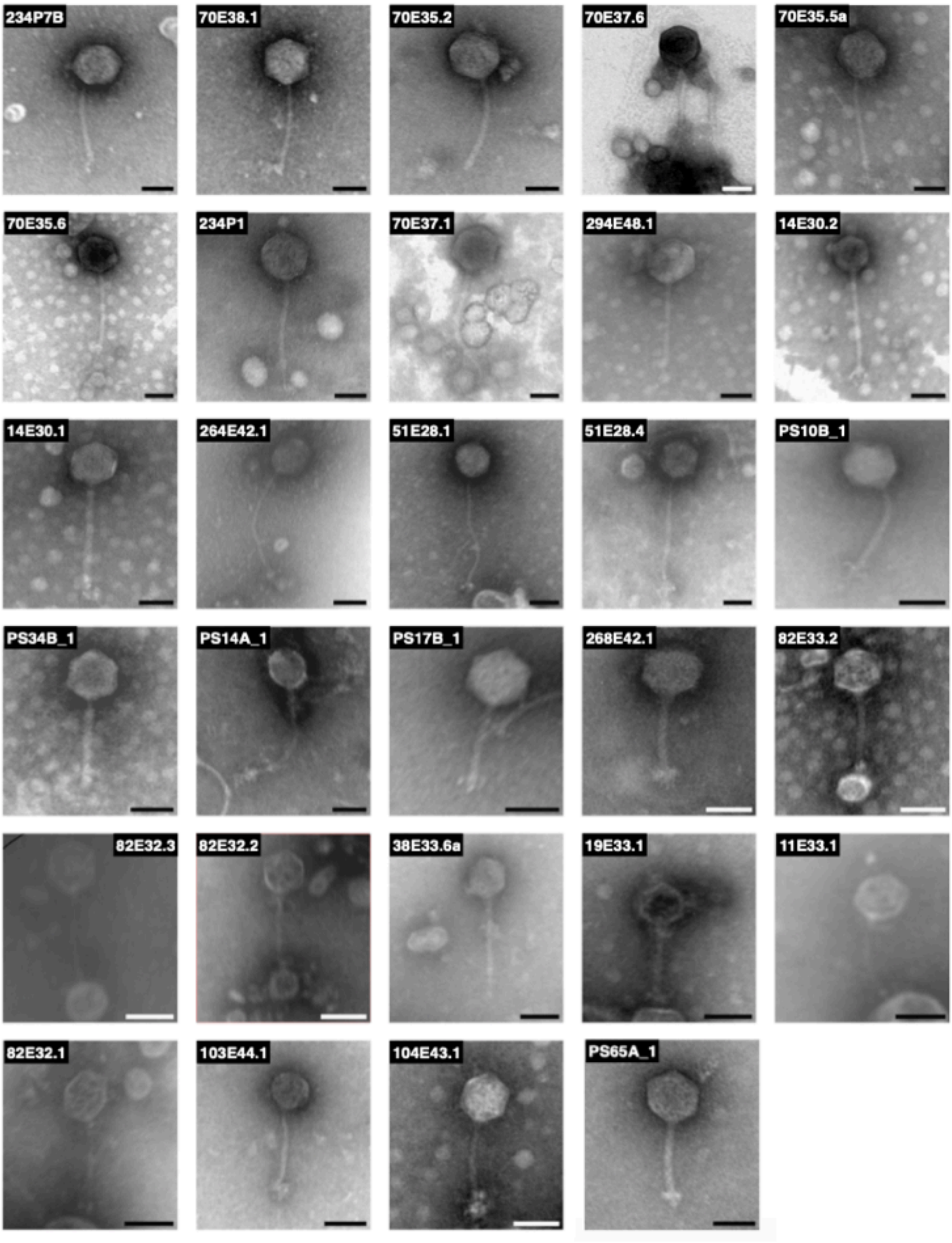


**Figure S7.** Transmission electron micrographs of 29 *Siphoviridae*. Scale bars represent 50nm

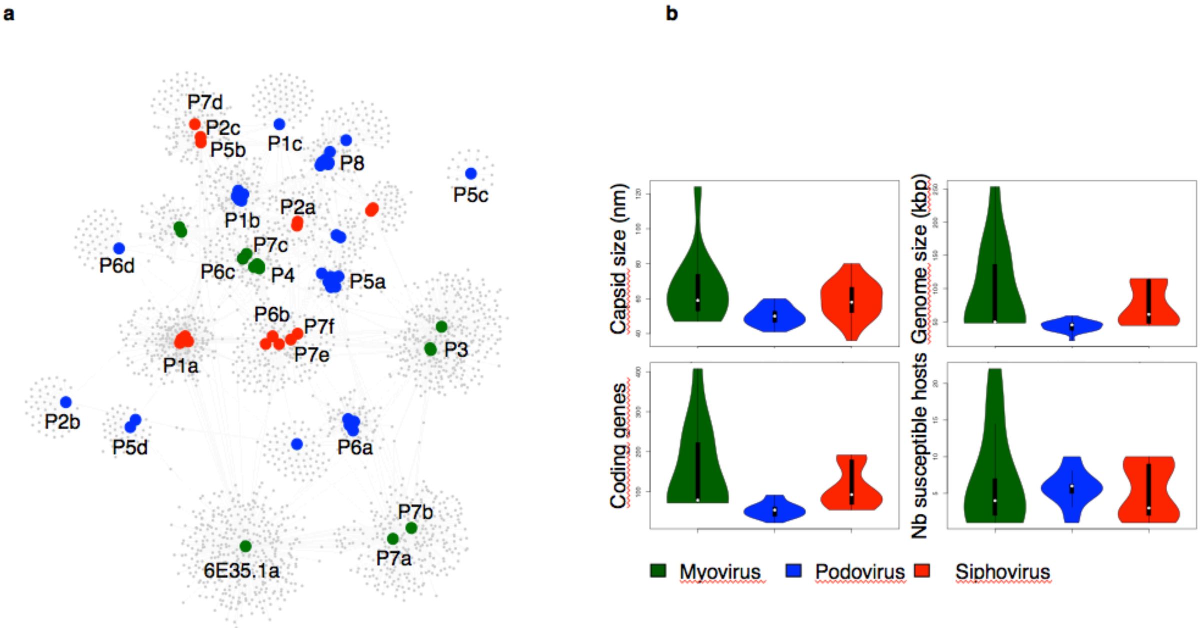


**Figure S8. General features of the sequenced phages. a,** Clustering of phage genomes using Cytoscape[19]. The network was integrated with 2,486 genes family from 76 phages and revealed clustering of phages with genomes (large circles) linked by common genes (small grey circles), %id aa>30% and >80% coverage. The color of each phage genome indicates the morphotype as in b. **b,** Capsid size, genome size and gene number differed significantly between morphotypes (capsid size: $F_{2,61} = 8.244$, $p < 0.001$, genome size: $F_{2,61} = 9.957$, $p < 0.001$, coding genes: $F_{2,61} = 12.1$, $p < 0.001$) while there was no difference in terms of susceptible vibrios between virus types ($F_{2,68} = 0.505$, $p < 0.606$). Podoviruses are smaller in capsid and genome size and contain fewer genes than both myoviruses and siphoviruses (Tukey’s HSD, all p values < 0.017), which showed no significant differences between each other for any measurements (Turkey’ HSD, all p values > 0.124).

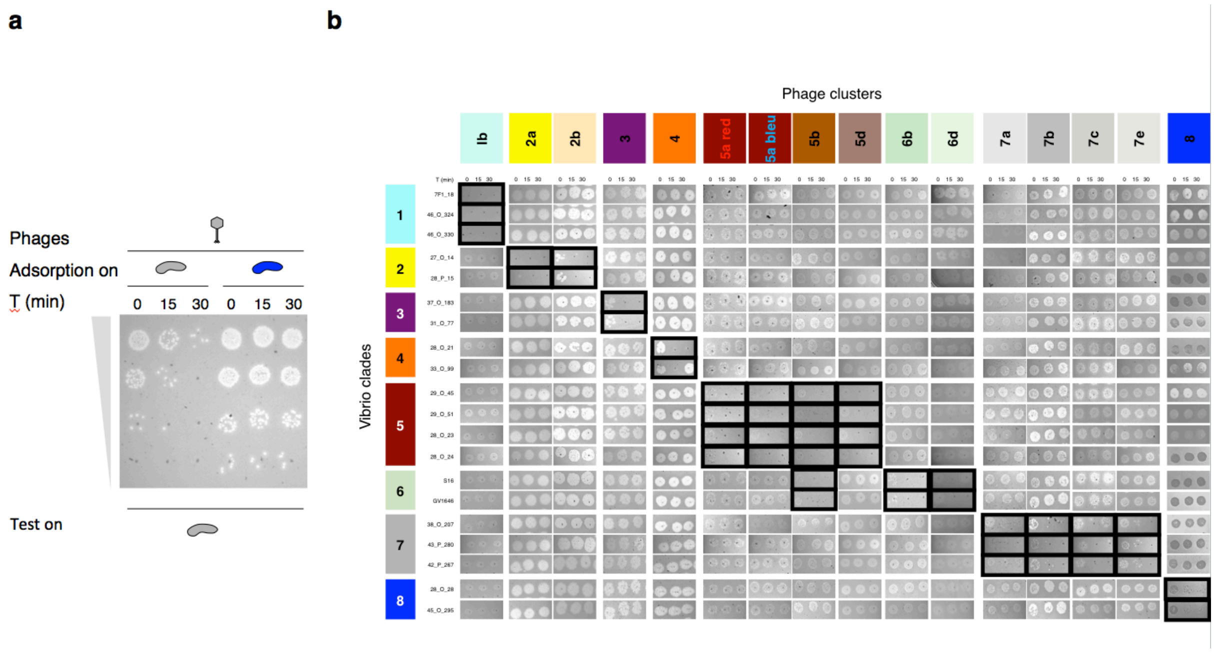


**Figure S9. Phage adsorption assays using representatives of the vibrio clades and phage clusters.** After allowing a fixed concentration of phages (MOI 0.01) to adsorb to a *V. crassostreae* strain, free phages that remained unattached were serially diluted and plated with sensitive hosts to quantify adsorption. All adsorption assays were performed twice. **a,** Example of the phage 268E42.1 from cluster P7e tested for adsorption on vibrio 43_P_280 and 28_O_28 from clade V7 and V8, respectively, and the serial dilution were plated on the sensitive vibrio 42_P_267. **b,** All 320 adsorption assays with positive results surrounded by bold.

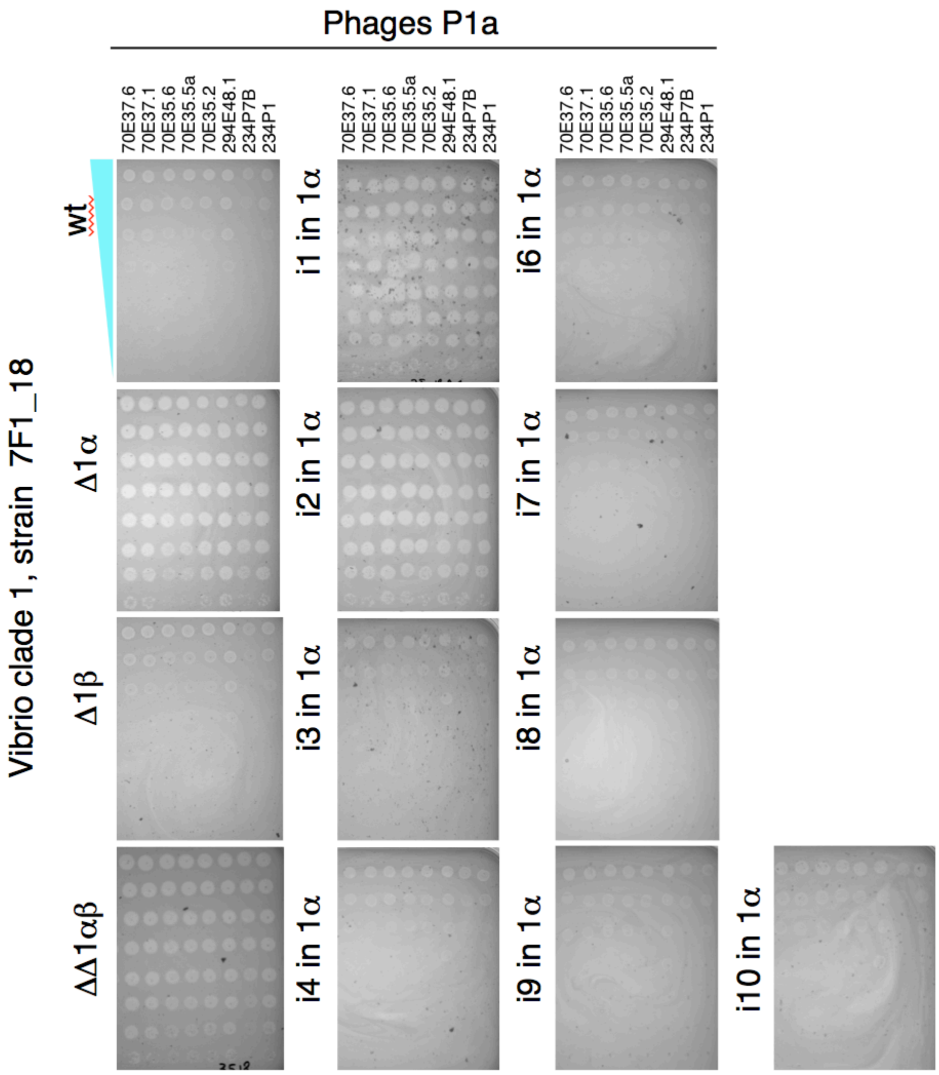


**Figure S10.** Changes in susceptibility to phage killing observed for phage-defense region deletions or gene inactivation in one strain (7F1_18) of vibrio clade V1 and phage from cluster P1a. Ten fold dilutions of phage from cluster P1a were plate on the strain 7F1_18 wild type and derivatives. All experiments were performed twice.

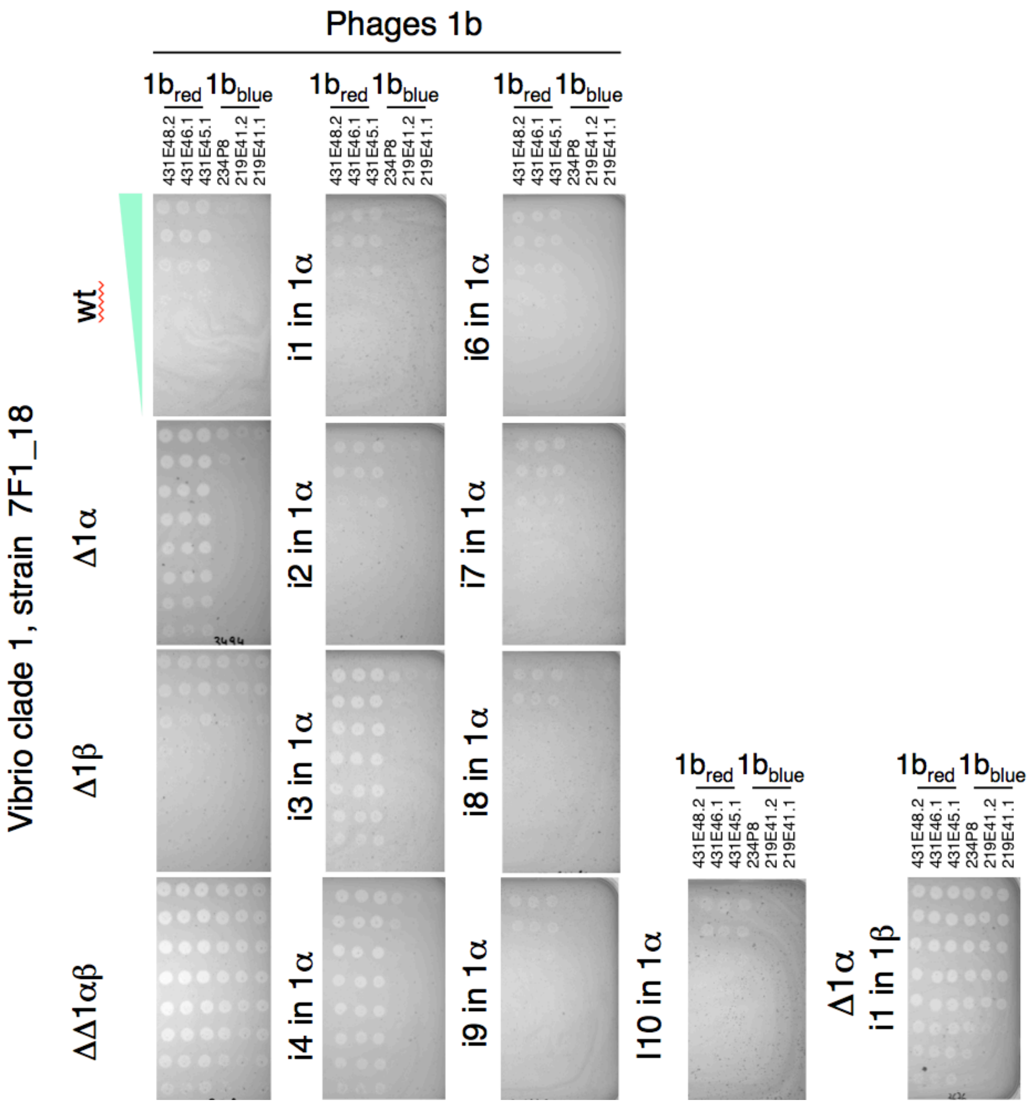


**Figure S11.** Changes in susceptibility to phage killing observed for phage-defense region deletions or gene inactivation in one strain (7F1_18) of vibrio clade V1. Tenfold dilutions of phages from cluster P1b were plate on the strain 7F1_18 wild type and derivatives. All experiments were performed twice.

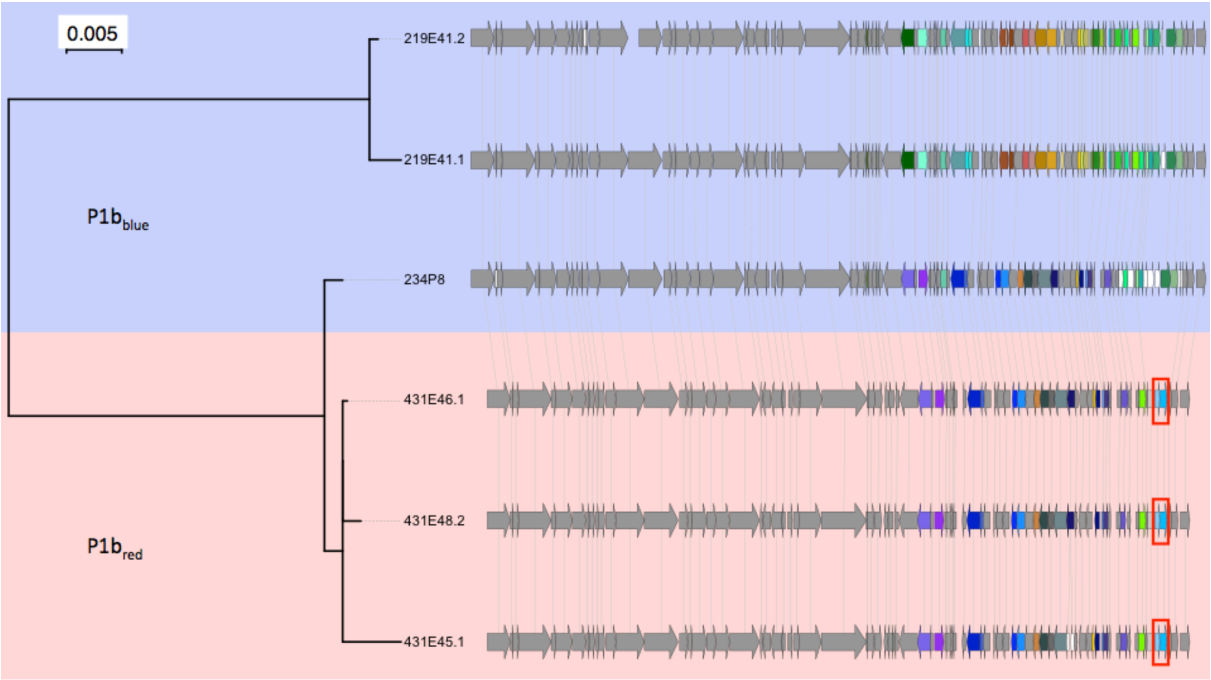


**Figure S12.** Genetic diversity within the phage cluster P1b. The core genome phylogeny demonstrates P1b$_{red}$ to a distinct clade. Gene synteny and content are overall conserved between all phages (in grey). Specific genes are colored with only two genes (framed in red) that were found in all P1b$_{red}$ phages.

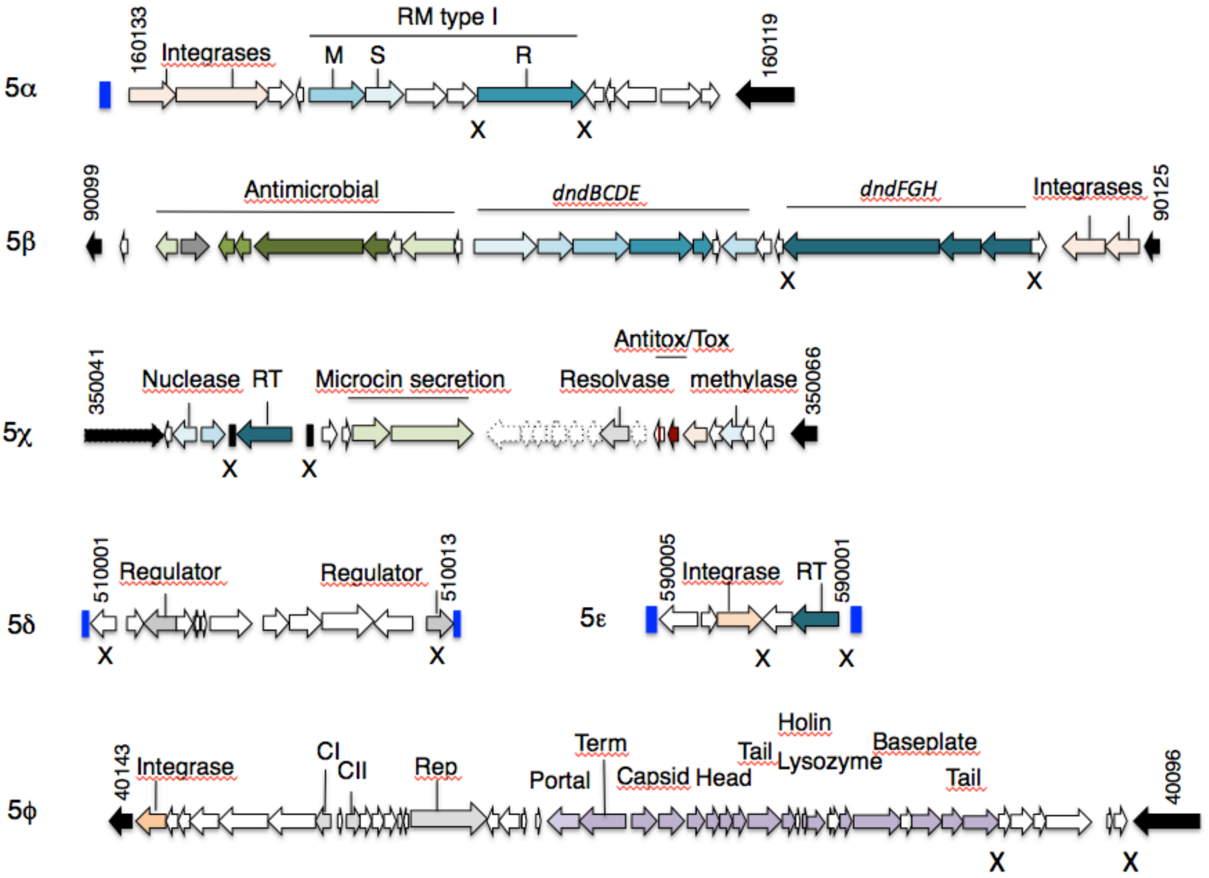


**Figure S13.** Gene diagrams of the genomic regions specific to strains $V5_{red}$ that are resistant to phage $P5a_{blue}$. X indicates the gene (s) that were deleted in each genomic region to explore their role for resistance.

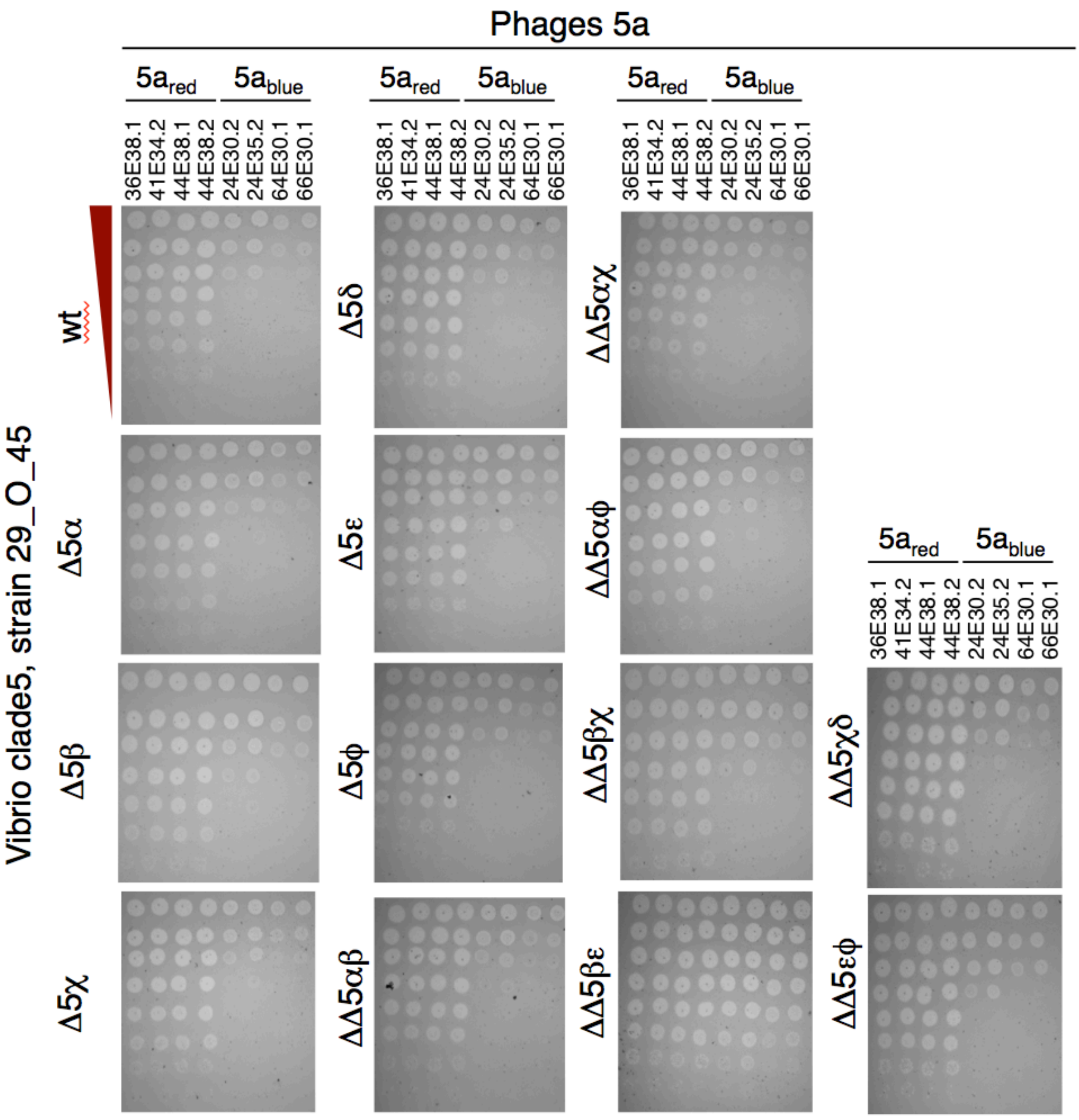


**Figure S14.** Changes in susceptibility to phage killing observed for phage-defense region deletions in one strain (29_O_45) of vibrio clade V5. Lawns of bacterial hosts with drop spots of a 1:10 dilution series of phages from cluster P5a. All experiments were performed twice.

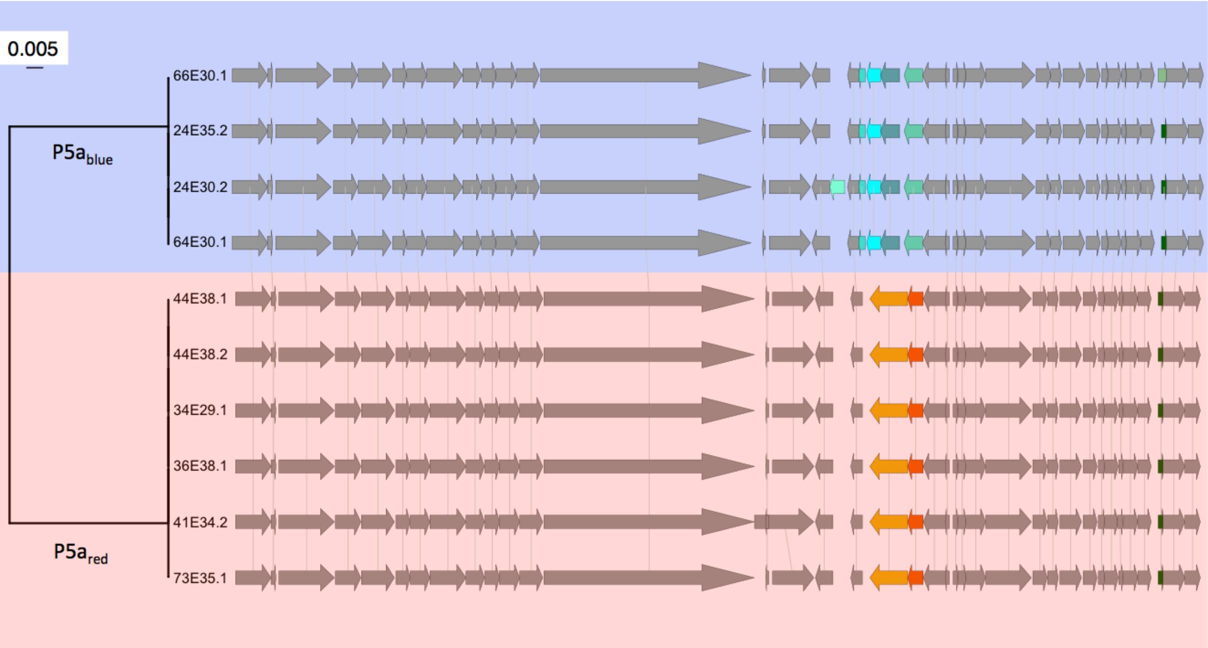


**Figure S15.** Genetic diversity within the phage cluster P5a. The core genome phylogeny demonstrates P5a$_{red}$ and P5a$_{blue}$ belong to distinct clades. Gene synteny and content are conserved between all P5a phages and 2 (orange, red) and 4 genes (blue, green) are specific to P5$_{red}$ and P5$_{blue}$ respectively.

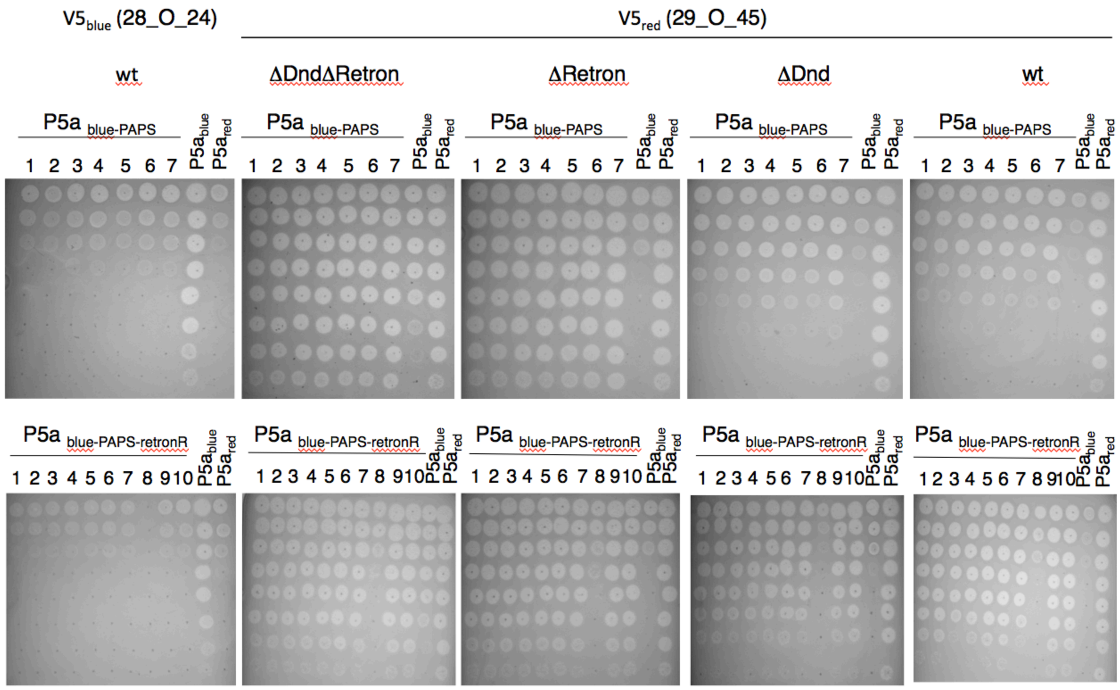


**Figure S16.** Changes in susceptibility to phage killing observed for phage-defense region deletions in one strain (29_O_45) of vibrio clade V5. Lawns of bacterial hosts with drop spots of a 1:10 dilution series of phages P5a$_{red}$, P5a$_{blue}$, and several isolates of P5a$_{blue}$ mutants. All experiments were performed twice.

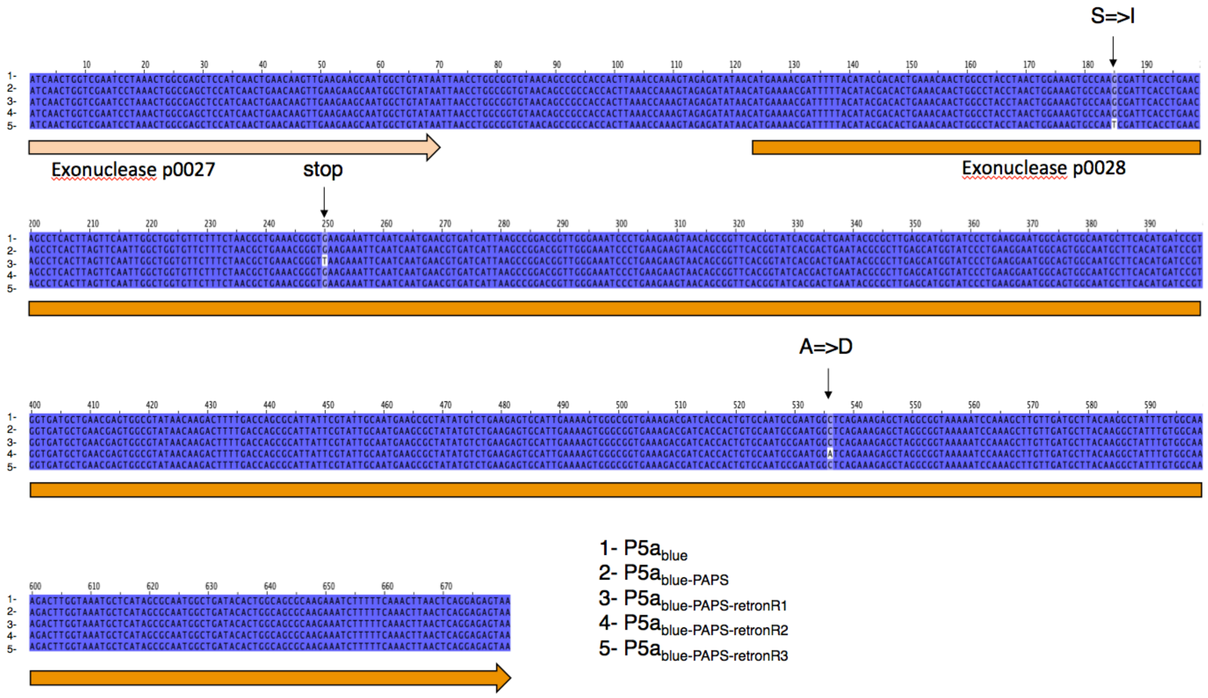


**Figure S17** Identification of the mutations potentially involved in retron escape. Sequencing the 5.7kb divergent regions of three retron-escaper phages revealed non-synonymous mutations or stop codon that distinguished the mutants from the ancestor, all localized in an exonuclease (p0028 in 66E30.1).

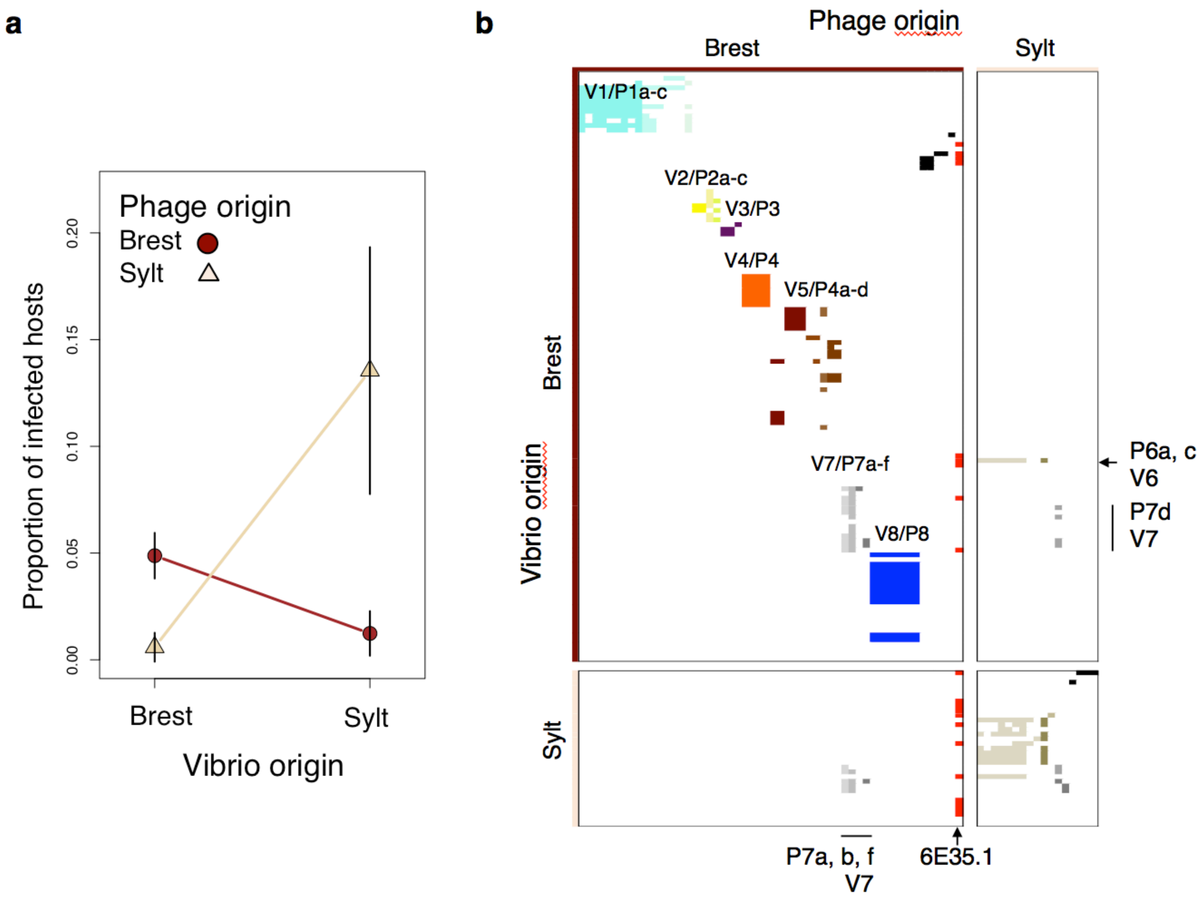


**Figure S18** Spatial phage-vibrio interactions. **a,** The all-by-all host range infection assay reveals that sympatric killing of vibrio by phages (Brest: 56 phages, 120 *Vibrio* isolates, Sylt: 17 phages, 33 Vibrio isolates) is more frequent than allopatric killing (Brest/Sylt) with the *Vibrio* origin * phage origin explaining most of the variation in the data (Analysis of Deviance based on Type III Wald chisquare tests, *Vibrio* origin Chi2 = 18.119, p = 2.1 e-05, phage origin Chi2 = 39.552, p = 3.2 e-10, *Vibrio* * phage origin chi2 = 161.063, p < 2.2e-16). **b,** distribution of phage cluster (P) and vibrio clade (V) in Sylt and Brest and interaction network.

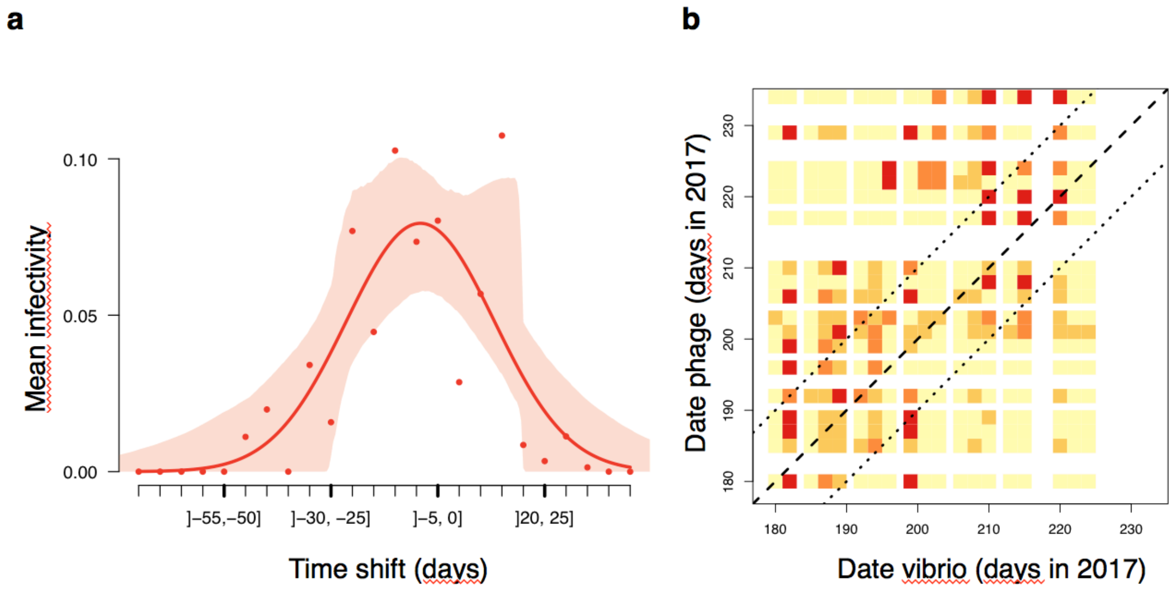


**Figure S19** Temporal pattern of adaptation at smaller time scales by using the intensive sampling carried out in Brest. **a,** Mean infectivity as a function of the time-shift in days between phages and vibrio using 1 to 15 *V. crassostreae* colonies per sampling event (median 3 colonies), and 1 to 47 phage strains (median 8 strains) in a total of 25,794 cross-inoculations. The points are the means, the curve and CI show the fit of a smooth function (proportional to the density of a skew-normal distribution) **b,** shows the mean infectivity as a function of the date of *V. crassostreae* and date of phages in cross-inoculation experiments. Yellow: mean infectivity 0, light orange: mean infectivity ]0, 0.1[, dark orange: mean infectivity [0.1, 0.2[, red: mean infectivity >=0.2. The diagonal (dashed line) shows the contemporaneous combinations, the sub-diagonals (dotted lines) the combinations shifted by ±10 days.
